# A multi-layer functional genomic analysis to understand noncoding genetic variation in lipids

**DOI:** 10.1101/2021.12.07.470215

**Authors:** Shweta Ramdas, Jonathan Judd, Sarah E Graham, Stavroula Kanoni, Yuxuan Wang, Ida Surakka, Brandon Wenz, Shoa L Clarke, Alessandra Chesi, Andrew Wells, Konain Fatima Bhatti, Sailaja Vedantam, Thomas W Winkler, Adam E Locke, Eirini Marouli, Greg JM Zajac, Kuan-Han H Wu, Ioanna Ntalla, Qin Hui, Derek Klarin, Austin T Hilliard, Zeyuan Wang, Chao Xue, Gudmar Thorleifsson, Anna Helgadottir, Daniel F Gudbjartsson, Hilma Holm, Isleifur Olafsson, Mi Yeong Hwang, Sohee Han, Masato Akiyama, Saori Sakaue, Chikashi Terao, Masahiro Kanai, Wei Zhou, Ben M Brumpton, Humaira Rasheed, Aki S Havulinna, Yogasudha Veturi, Jennifer Allen Pacheco, Elisabeth A Rosenthal, Todd Lingren, QiPing Feng, Iftikhar J. Kullo, Akira Narita, Jun Takayama, Hilary C Martin, Karen A Hunt, Bhavi Trivedi, Jeffrey Haessler, Franco Giulianini, Yuki Bradford, Jason E Miller, Archie Campbell, Kuang Lin, Iona Y Millwood, Asif Rasheed, George Hindy, Jessica D Faul, Wei Zhao, David R Weir, Constance Turman, Hongyan Huang, Mariaelisa Graff, Ananyo Choudhury, Dhriti Sengupta, Anubha Mahajan, Michael R Brown, Weihua Zhang, Ketian Yu, Ellen M Schmidt, Anita Pandit, Stefan Gustafsson, Xianyong Yin, Jian’an Luan, Jing-Hua Zhao, Fumihiko Matsuda, Hye-Mi Jang, Kyungheon Yoon, Carolina Medina-Gomez, Achilleas Pitsillides, Jouke Jan Hottenga, Andrew R Wood, Yingji Ji, Zishan Gao, Simon Haworth, Ruth E Mitchell, Jin Fang Chai, Mette Aadahl, Anne A Bjerregaard, Jie Yao, Ani Manichaikul, Wen-Jane, A Chao, Helen R Warren, Julia Ramirez, Jette Bork-Jensen, Line L Kårhus, Anuj Goel, Maria Sabater-Lleal, Raymond Noordam, Pala Mauro, Floris Matteo, Aaron F McDaid, Pedro Marques-Vidal, Matthias Wielscher, Stella Trompet, Naveed Sattar, Line T Møllehave, Matthias Munz, Lingyao Zeng, Jianfeng Huang, Bin Yang, Alaitz Poveda, Azra Kurbasic, Sebastian Schönherr, Lukas Forer, Markus Scholz, Tessel E. Galesloot, Jonathan P. Bradfield, Sanni E Ruotsalainen, E Warwick Daw, Joseph M Zmuda, Jonathan S Mitchell, Christian Fuchsberger, Henry Christensen, Jennifer A Brody, Phuong Le, Mary F Feitosa, Mary K Wojczynski, Daiane Hemerich, Michael Preuss, Massimo Mangino, Paraskevi Christofidou, Niek Verweij, Jan W Benjamins, Jorgen Engmann, Tsao L. Noah, Anurag Verma, Roderick C Slieker, Ken Sin Lo, Nuno R Zilhao, Marcus E Kleber, Graciela E Delgado, Shaofeng Huo, Daisuke D Ikeda, Hiroyuki Iha, Jian Yang, Jun Liu, Ayşe Demirkan, Hampton L Leonard, Jonathan Marten, Carina Emmel, Börge Schmidt, Laura J Smyth, Marisa Cañadas-Garre, Chaolong Wang, Masahiro Nakatochi, Andrew Wong, Nina Hutri-Kähönen, Xueling Sim, Rui Xia, Alicia Huerta-Chagoya, Juan Carlos Fernandez-Lopez, Valeriya Lyssenko, Suraj S Nongmaithem, Alagu Sankareswaran, Marguerite R Irvin, Christopher Oldmeadow, Han-Na Kim, Seungho Ryu, Paul RHJ Timmers, Liubov Arbeeva, Rajkumar Dorajoo, Leslie A Lange, Gauri Prasad, Laura Lorés-Motta, Marc Pauper, Jirong Long, Xiaohui Li, Elizabeth Theusch, Fumihiko Takeuchi, Cassandra N Spracklen, Anu Loukola, Sailalitha Bollepalli, Sophie C Warner, Ya Xing Wang, Wen B. Wei, Teresa Nutile, Daniela Ruggiero, Yun Ju Sung, Shufeng Chen, Fangchao Liu, Jingyun Yang, Katherine A Kentistou, Bernhard Banas, Anna Morgan, Karina Meidtner, Lawrence F Bielak, Jennifer A Smith, Prashantha Hebbar, Aliki-Eleni Farmaki, Edith Hofer, Maoxuan Lin, Maria Pina Concas, Simona Vaccargiu, Peter J van der Most, Niina Pitkänen, Brian E Cade, Sander W. van der Laan, Kumaraswamy Naidu Chitrala, Stefan Weiss, Amy R Bentley, Ayo P Doumatey, Adebowale A Adeyemo, Jong Young Lee, Eva RB Petersen, Aneta A Nielsen, Hyeok Sun Choi, Maria Nethander, Sandra Freitag-Wolf, Lorraine Southam, Nigel W Rayner, Carol A Wang, Shih-Yi Lin, Jun-Sing Wang, Christian Couture, Leo-Pekka Lyytikäinen, Kjell Nikus, Gabriel Cuellar-Partida, Henrik Vestergaard, Bertha Hidalgo, Olga Giannakopoulou, Qiuyin Cai, Morgan O Obura, Jessica van Setten, Karen Y. He, Hua Tang, Natalie Terzikhan, Jae Hun Shin, Rebecca D Jackson, Alexander P Reiner, Lisa Warsinger Martin, Zhengming Chen, Liming Li, Takahisa Kawaguchi, Joachim Thiery, Joshua C Bis, Lenore J Launer, Huaixing Li, Mike A Nalls, Olli T Raitakari, Sahoko Ichihara, Sarah H Wild, Christopher P Nelson, Harry Campbell, Susanne Jäger, Toru Nabika, Fahd Al-Mulla, Harri Niinikoski, Peter S Braund, Ivana Kolcic, Peter Kovacs, Tota Giardoglou, Tomohiro Katsuya, Dominique de Kleijn, Gert J. de Borst, Eung Kweon Kim, Hieab H.H. Adams, M. Arfan Ikram, Xiaofeng Zhu, Folkert W Asselbergs, Adriaan O Kraaijeveld, Joline WJ Beulens, Xiao-Ou Shu, Loukianos S Rallidis, Oluf Pedersen, Torben Hansen, Paul Mitchell, Alex W Hewitt, Mika Kähönen, Louis Pérusse, Claude Bouchard, Anke Tönjes, Yii-Der Ida Chen, Craig E Pennell, Trevor A Mori, Wolfgang Lieb, Andre Franke, Claes Ohlsson, Dan Mellström, Yoon Shin Cho, Hyejin Lee, Jian-Min Yuan, Woon-Puay Koh, Sang Youl Rhee, Jeong-Taek Woo, Iris M Heid, Klaus J Stark, Martina E Zimmermann, Henry Völzke, Georg Homuth, Michele K Evans, Alan B Zonderman, Ozren Polasek, Gerard Pasterkamp, Imo E Hoefer, Susan Redline, Katja Pahkala, Albertine J Oldehinkel, Harold Snieder, Ginevra Biino, Reinhold Schmidt, Helena Schmidt, Stefania Bandinelli, George Dedoussis, Thangavel Alphonse Thanaraj, Patricia A Peyser, Norihiro Kato, Matthias B Schulze, Giorgia Girotto, Carsten A Böger, Bettina Jung, Peter K Joshi, David A Bennett, Philip L De Jager, Xiangfeng Lu, Vasiliki Mamakou, Morris Brown, Mark J Caulfield, Patricia B Munroe, Xiuqing Guo, Marina Ciullo, Jost B. Jonas, Nilesh J Samani, Jaakko Kaprio, Päivi Pajukanta, Teresa Tusié-Luna, Carlos A Aguilar-Salinas, Linda S Adair, Sonny Augustin Bechayda, H. Janaka de Silva, Ananda R Wickremasinghe, Ronald M Krauss, Jer-Yuarn Wu, Wei Zheng, Anneke I den Hollander, Dwaipayan Bharadwaj, Adolfo Correa, James G Wilson, Lars Lind, Chew-Kiat Heng, Amanda E Nelson, Yvonne M Golightly, James F Wilson, Brenda Penninx, Hyung-Lae Kim, John Attia, Rodney J Scott, D C Rao, Donna K Arnett, Mark Walker, Laura J Scott, Heikki A Koistinen, Giriraj R Chandak, Josep M Mercader, Teresa Tusie-Luna, Carlos Aguilar-Salinas, Clicerio Gonzalez Villalpando, Lorena Orozco, Myriam Fornage, E Shyong Tai, Rob M van Dam, Terho Lehtimäki, Nish Chaturvedi, Mitsuhiro Yokota, Jianjun Liu, Dermot F Reilly, Amy Jayne McKnight, Frank Kee, Karl-Heinz Jöckel, Mark I McCarthy, Colin NA Palmer, Veronique Vitart, Caroline Hayward, Eleanor Simonsick, Cornelia M van Duijn, Zi-Bing Jin, Fan Lu, Haretsugu Hishigaki, Xu Lin, Winfried März, Vilmundur Gudnason, Jean-Claude Tardif, Guillaume Lettre, Leen M t Hart, Petra JM Elders, Daniel J Rader, Scott M Damrauer, Meena Kumari, Mika Kivimaki, Pim van der Harst, Tim D Spector, Ruth J.F. Loos, Michael A Province, Esteban J Parra, Miguel Cruz, Bruce M Psaty, Ivan Brandslund, Peter P Pramstaller, Charles N Rotimi, Kaare Christensen, Samuli Ripatti, Elisabeth Widén, Hakon Hakonarson, Struan F.A. Grant, Lambertus ALM Kiemeney, Jacqueline de Graaf, Markus Loeffler, Florian Kronenberg, Dongfeng Gu, Jeanette Erdmann, Heribert Schunkert, Paul W Franks, Allan Linneberg, J. Wouter Jukema, Amit V Khera, Minna Männikkö, Marjo-Riitta Jarvelin, Zoltan Kutalik, Cucca Francesco, Dennis O Mook-Kanamori, Ko Willems van Dijk, Hugh Watkins, David P Strachan, Niels Grarup, Peter Sever, Neil Poulter, Wayne Huey-Herng Sheu, Jerome I Rotter, Thomas M Dantoft, Fredrik Karpe, Matt J Neville, Nicholas J Timpson, Ching-Yu Cheng, Tien-Yin Wong, Chiea Chuen Khor, Hengtong Li, Charumathi Sabanayagam, Annette Peters, Christian Gieger, Andrew T Hattersley, Nancy L Pedersen, Patrik KE Magnusson, Dorret I Boomsma, Eco JC de Geus, L Adrienne Cupples, Joyce B.J. van Meurs, Arfan Ikram, Mohsen Ghanbari, Penny Gordon-Larsen, Wei Huang, Young Jin Kim, Yasuharu Tabara, Nicholas J Wareham, Claudia Langenberg, Eleftheria Zeggini, Jaakko Tuomilehto, Johanna Kuusisto, Markku Laakso, Erik Ingelsson, Goncalo Abecasis, John C Chambers, Jaspal S Kooner, Paul S de Vries, Alanna C Morrison, Scott Hazelhurst, Michèle Ramsay, Kari E. North, Martha Daviglus, Peter Kraft, Nicholas G Martin, John B Whitfield, Shahid Abbas, Danish Saleheen, Robin G Walters, Michael V Holmes, Corri Black, Blair H Smith, Aris Baras, Anne E Justice, Julie E Buring, Paul M Ridker, Daniel I Chasman, Charles Kooperberg, Gen Tamiya, Masayuki Yamamoto, David A van Heel, Richard C Trembath, Wei-Qi Wei, Gail P Jarvik, Bahram Namjou, M. Geoffrey Hayes, Marylyn D Ritchie, Pekka Jousilahti, Veikko Salomaa, Kristian Hveem, Bjørn Olav Åsvold, Michiaki Kubo, Yoichiro Kamatani, Yukinori Okada, Yoshinori Murakami, Bong-Jo Kim, Unnur Thorsteinsdottir, Kari Stefansson, Jifeng Zhang, Y Eugene Chen, Yuk-Lam Ho, Julie A Lynch, Daniel Rader, Philip S Tsao, Kyong-Mi Chang, Kelly Cho, Christopher J O’Donnell, John M Gaziano, Peter Wilson, Karen L Mohlke, Timothy M Frayling, Joel N Hirschhorn, Sekar Kathiresan, Michael Boehnke, Million Veterans Program, Global Lipids Genetics Consortium, Struan Grant, Pradeep Natarajan, Yan V Sun, Andrew P Morris, Panos Deloukas, Gina Peloso, Themistocles L Assimes, Cristen J Willer, Xiang Zhu, Christopher D Brown

**Author notes:** Corresponding Authors: Xiang Zhu, PhD, Department of Statistics, Huck Institutes of the Life Sciences, The Pennsylvania State University, University Park, PA 16802, Christopher D Brown, PhD, Department of Genetics, Perelman School of Medicine, University of Pennsylvania, Philadelphia PA 19104.

## Abstract

A major challenge of genome-wide association studies (GWAS) is to translate phenotypic associations into biological insights. Here, we integrate a large GWAS on blood lipids involving 1.6 million individuals from five ancestries with a wide array of functional genomic datasets to discover regulatory mechanisms underlying lipid associations. We first prioritize lipid-associated genes with expression quantitative trait locus (eQTL) colocalizations, and then add chromatin interaction data to narrow the search for functional genes. Polygenic enrichment analysis across 697 annotations from a host of tissues and cell types confirms the central role of the liver in lipid levels, and highlights the selective enrichment of adipose-specific chromatin marks in high-density lipoprotein cholesterol and triglycerides. Overlapping transcription factor (TF) binding sites with lipid-associated loci identifies TFs relevant in lipid biology. In addition, we present an integrative framework to prioritize causal variants at GWAS loci, producing a comprehensive list of candidate causal genes and variants with multiple layers of functional evidence. Two prioritized genes, *CREBRF* and *RRBP1*, show convergent evidence across functional datasets supporting their roles in lipid biology.

## Introduction

Most GWAS findings have not directly led to mechanistic interpretations, largely because 90% of GWAS associations map to non-coding sequences ^1,2^. Mechanistic interpretations in GWAS have proven challenging because the strongest signals identified in GWAS typically contain many variants in strong linkage disequilibrium (LD) ^3^ and functional mechanisms including genes of action are often not clear from GWAS data alone ^4,5^.

Linking trait-associated variants to genome function has emerged as a promising model for mechanistic interpretation of non-coding findings in GWAS. This ‘variant-to-function’ model is premised on recent observations that non-coding variants often affect a trait of interest through the regulation of genes and processes in trait-relevant cell types or tissues ^2,6^. Implementing this functional model in GWAS has become more feasible as large-scale functional genomic resources, such as epigenomic ^7^ and transcriptomic ^8^ catalogues, have been systematically generated across a wide range of human cell types and tissues. The integration of functional genomics with GWAS has identified regulatory mechanisms in variants associated with some flagship disorders such as obesity ^9^ and schizophrenia ^10^, yielding important functional insights into the genetic architecture of human complex traits.

The history of the human genetics of lipids mirrors the successes and challenges of GWAS. Increasing sample size and genetic diversity has significantly boosted the power of discovery: the first lipid GWAS in 2008 with 8,816 European-descent individuals identified 29 lipid-associated loci^11^; the latest study of 1.6 million individuals across five ancestries ^12^ found 941. Despite the dramatic increase in the number of associations, our biological understanding of many of these genetic discoveries remains limited. The causal gene has been confidently assigned at only a small fraction of these loci ^2^, and the regulatory mechanism connecting variant to phenotype has been conclusively characterized only for a handful of genes ^5^. Furthermore, systematic mapping of lipid-associated variants to their biological functions has been missing in the literature at the time of this study.

Here we conduct a genome-scale integrative analysis on the largest GWAS to-date of five lipid phenotypes (LDL, or low density lipoprotein; HDL, or high density lipoprotein; TC, or total cholesterol; nonHDL, or non-high density lipoprotein; and TG, or triglycerides) involving 1.65 million individuals from five ancestries ^12^. Combining the lipid GWAS with a wide array of functional genomic resources in diverse human tissues and cell types, we identify regulatory mechanisms of noncoding genetic variation in lipids with a full suite of computational approaches. Further, we develop a generalizable framework to understand how tissue-specific gene regulation can explain GWAS findings, and demonstrate its real-world value on lipid-associated loci.

## Material and Methods

### GWAS

We performed GWAS for five blood lipid traits (LDL, HDL, TC, TG, and nonHDL) in 1.65 million individuals from five ancestry groups ^12^(African and African-admixed, East Asian, European, Hispanic, South Asian) at 91 million variants imputed primarily from the Haplotype Reference Consortium ^13^ or 1000 Genomes Phase 3 ^14^. The individual GWAS and meta-analyses were performed using the hg19 version of the human reference genome. We used MR-MEGA ^15^ for meta-analysis across cohorts.

We defined ‘sentinel variants’ as lead variants representing independent trait-associated loci in the genome. These windows are the greater of 500kb or 0.25cM around the sentinel variant; genetic distances were defined using reference maps from HapMap 3 ^16^. We performed a second round of conditional analysis, conditioning on the sentinel variants to identify and remove any significant windows that are shadow signals (or dependent on) of a neighboring locus to enforce independence of associated loci.

### Colocalization with eQTLs

We performed statistical colocalization of lipid GWAS with eQTLs obtained from GTEx v8 across 49 tissues ^8^. For each of the five lipid traits, we used the same sentinel variants defined in the previous section to represent approximately independent GWAS-associated windows (also removing shadow signals as described before).

For each such window, we ran eQTL colocalization with GTEx v8 single-tissue cis-eQTL summary statistics ^8^. For each of 49 GTEx tissues, we first identified all genes within 1Mb of the sentinel SNP, and then restricted analysis to those genes with significant eQTLs (i.e., ‘eGenes’ as defined by GTEx) in that tissue (FDR < 0.05). We used the R package ‘coloc’ (run on R version 3.4.3, coloc version 3.2.1) ^17^ with default parameters to run colocalization between the GWAS signal and the eQTL signal for each of these cis-eGenes, using as input those SNPs in the defined window (greater than 500kb or 0.25cM on either side of the lead variant), i.e. all SNPs present in both datasets. eQTL summary statistics were in GRCh38, so we first lifted over the GWAS summary statistics (in hg19) to GRCh38 using liftOver ^18^. As in previous studies ^19^, we used a colocalization posterior probability of (PP3+PP4) > 0.8 to identify loci with enough colocalization power, and PP4/PP3 > 0.9 to define those loci that show significant colocalization, where PP4 represents posterior probability of a single shared signal, and PP3 represents posterior probability of two unique signals in the GWAS and eQTL datasets.

### Overlap with promoter Capture-C data

We used four promoter-focused Capture-C (henceforth Capture-C) datasets from three human cell/tissue types to capture physical interactions between gene promoters and their regulatory elements. We used three biological replicates of HepG2 liver carcinoma cells ^20^, another HepG2 dataset described in Selvarajan et al ^21^, hepatocyte-like cells (HLC) produced by differentiating three biological replicates of iPSCs (which in turn were generated from peripheral blood mononuclear cells using a previously published protocol ^22^), and an adipose dataset obtained from Pan et al ^23^ that was produced using primary human white adipocytes.

The detailed protocol to prepare HepG2 or HLC cells for the Capture-C experiment is described in Chesi et al^20^. Briefly, for each dataset, 10 million cells were used for promoter Capture-C library generation. Custom capture baits were designed using an Agilent SureSelect library design targeting both ends of DpnII restriction fragments encompassing promoters (including alternative promoters) of all human coding genes, noncoding RNA, antisense RNA, snRNA, miRNA, snoRNA, and lincRNA transcripts, totaling 36,691 RNA baited fragments. Each library was then sequenced on an Illumina HiSeq 4000 (HepG2) or Illumina NovoSeq (HLC), generating 1.6 billion read pairs per sample (50 base pair read length.) We used HiCUP v0.7.2 ^24^ to process the raw FastQ files into loop calls and CHiCAGO v1.6.0 ^24,25^ to define significant looping interactions; we defined a CHiCAGO score of 5 as significant, as specified in the default parameters.

Starting with Capture-C maps processed as described above, we re-annotated the baits to gene IDs from Gencode v19 ^26^ to ensure uniformity of gene annotations with the rest of our pipeline. For each bait, we identified any gene whose transcription start site (TSS) from any transcript in Gencode v19 was within 175 base pair distance from the bait (to account for differing bait designs for external datasets which may not directly overlap the canonical TSS). We filtered all datasets to only include interactions in which the interacting end was not another bait. Enrichment with colocalized genes was robust to our choice of distance between bait and gene (enrichment with eQTL colocalized genes ranging from 2.94-2.96 for bait distances from 0-350 base pairs).

To identify genetic variants associated with any of the five lipid traits that physically interact with locations in the genome, we used the R package ‘Genomic Ranges’ version 1.30.3 ^27^ to find overlap between credible sets for each trait’s GWAS and the previously annotated promoter Capture-C data; we refer to these as Capture-C/GWAS interactions. Each credible set was defined as the set of variants with a 95% posterior probability of being the causal variant. For all individual variants within all GWAS-associated loci for the five lipid traits, we identified which variants overlapped any interacting end of the four previously annotated promoter Capture-C data.

### Presence of gene-variant pairs in same topologically associated domains

To estimate the frequency of colocalized gene-sentinel pairs in the same topologically associated domain (TAD), we used publicly-available TADs from human liver ^28^. We compared the number of colocalizations with the sentinel variant and colocalized gene in the same TAD divided by all colocalizations in which the sentinel variant lies in a TAD. To test if this ratio was statistically significant, we generated random TAD boundaries using ‘bedtools shuffle’ 1000 times, and calculated the same ratio for these randomly-generated TAD boundaries.

### Pathway enrichment

We used ClusterProfiler v3.6.0 ^29^ to look for pathways over-represented in each gene list: genes with eQTL colocalization and genes interacting with variants in GWAS credible sets. We used the enrichKEGG function to look for pathway enrichment in KEGG pathways (using the latest version of the KEGG database ^30^). We first re-mapped gencode IDs to gene symbols using the Gencode v24 annotation and then used the biomaRt R package v2.34.2 ^31^ to convert gene symbols to Entrez IDs. We ran enrichKEGG to identify enriched pathways significant at a Benjamini-Hochberg threshold of 0.05.

### Enrichment in known lipid-associated genes

We calculated enrichment odds ratio of genes identified in our analysis with three known sets of lipid-associated genes using the Fisher’s exact test (R function ‘fisher.test’). First, we identified a list of 33 Mendelian genes from ClinVar ^32^ with lipidemia-associated ICD10 codes (E78). Second, we used the set of genes identified from a transcriptome-wide association study (TWAS) on the same GWAS and GTEx v8 summary statistics using the S-PrediXcan software ^33^ default setup. Third, we used 35 genes with rare-coding variants associated with lipid levels ^34^.

### Stratified LD score regression

We used LDSC version 1.0.1 ^35^ to estimate the enrichment of heritability using GWAS summary statistics in different epigenetic and transcriptomic annotations, including gene expression, chromatin marks and TF binding sites. The gene expression and chromatin mark annotations and the corresponding LD scores were provided as ‘Multitissuegeneexpr1000Gv3’ and ‘Multitissuechromatin1000Gv3’ databases in LDSC software. The TF binding site annotations were extracted from ChIP-seq data of 161 TFs from ENCODE, and their LD scores were estimated from 1000 Genomes Phase 3 European samples using ‘ldsc.py --l2’. We first converted the summary statistics for each phenotype to LDSC-formatted summary statistics using ‘munge_sumstats.py’. Second, we ran ‘ldsc.py’ using the baseline_v1.2 baseline model on each annotation to estimate enrichment of heritability. For primary analyses, we used multi-population GWAS summary statistics and LD scores estimated from 1000 Genomes Phase 3 European samples. For secondary analyses on East Asian GWAS alone, we obtained EAS-specific LD scores for the same epigenomic annotations ^36^.

### GREGOR analysis

We used GREGOR ^37^ to estimate enrichment of sentinel variants for each lipid phenotype in TF binding sites for 161 TFs from ENCODE compared to a null distribution of variants matched for allele frequency. We ran GREGOR with default parameters, specifying 0.8 as the R^2^ threshold, window size of 1Mb, and ‘EUR’ as the population. Annotations with FDR-adjusted P-value < 0.05 were considered significant.

### Enrichment in single-cell expression data

We overlapped our list of colocalized genes with publicly available single-cell RNA-sequencing data of 8,444 cells from liver ^38^ and 38,408 cells from adipose (Web resources) in humans. For both datasets, we downloaded normalized TPM data and existing tSNE cluster annotations for each cell. For each cluster, we defined median expression for each gene across all cells in that cluster. Then for each cluster, we calculated the enrichment P-value for our list of colocalized genes using the ‘fgsea’ R package v1.4.1, which looks for overrepresentation of our gene list in ranked genes for each cluster ^39^, implemented in R 3.4.3.

## Results

We systematically integrated lipid GWAS results ^12^ with multiple layers of functional genomic data from diverse tissues and cell types to understand regulatory mechanisms at lipid-associated loci (Figure 1). Specifically, we overlaid GWAS loci with eQTL and chromatin-chromatin interactions to identify causal genes. We assessed polygenic enrichments of tissue-specific histone marks to prioritize relevant tissues and examined GWAS loci at transcription factor (TF) binding sites to detect lipid-relevant TFs. Finally, we combined all these layers to prioritize functional variants at GWAS loci, providing a holistic view of gene regulation at lipid loci in relevant tissue and cell types.

**Figure 1:**
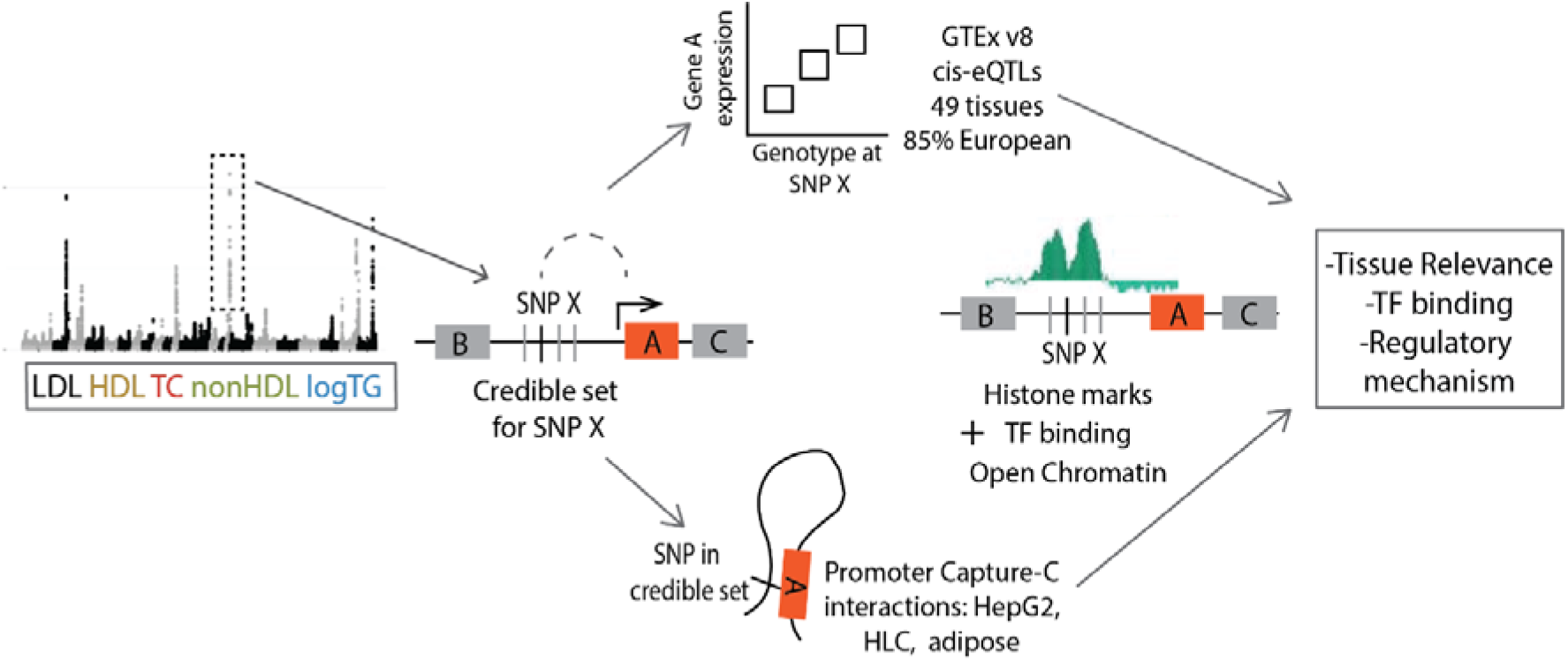
Schematic overview of the multi-layer functional genomic analysis. We first integrate GWAS summary statistics for five lipid phenotypes with eQTL and chromatin interaction data to identify potential genes mediating the GWAS association, and then incorporate epigenomic annotations to identify regulatory mechanisms at these loci. For any lead variant ‘X’, A, B, and C represent nearby eGenes, and SNPs around SNP X represent variants in the credible set.

### Colocalization with eQTLs identifies candidate lipid-relevant genes

First, we identified shared association signals between lipid levels and expression of nearby genes, since most GWAS signals are presumed to influence complex traits through impact on gene expression ^40^. To do so, we tested for colocalization of each of the 1,750 significant lipid GWAS association signals across the five traits examined with significant cis-eQTL data across 49 human tissues from the GTEx consortium ^8^. Here, we defined significant GWAS signals as 1,750 loci reaching genome-wide significance and corrected for shadow signals (Methods) in our multi-population meta-analysis for at least one of five lipid traits.

Second, we restricted our analysis to those loci likely mediated through regulatory mechanisms as opposed to coding variation. In particular, we excluded all loci with credible sets containing at least one missense variant (369 of 1,750 loci, 21% of credible sets). Of the remaining 1,381 GWAS loci, 696 significantly colocalized with eQTLs (the ratio of posterior probability of a shared signal to the posterior probability of two signals being > 0.9 ^19^; Methods) in at least one of 49 tissues for at least one lipid phenotype. This resulted in 1,076 colocalized eGenes ranging from 1 to 16 genes per locus (Figure 2A; Table S1). Since with eQTL data alone it is difficult to disentangle a single functional gene from multiple functional (and likely coregulated) genes at a locus ^41^ we performed all downstream analyses with all 1,076 colocalized genes, to further prioritize functional genes at loci with multiple eGenes.

**Figure 2:**
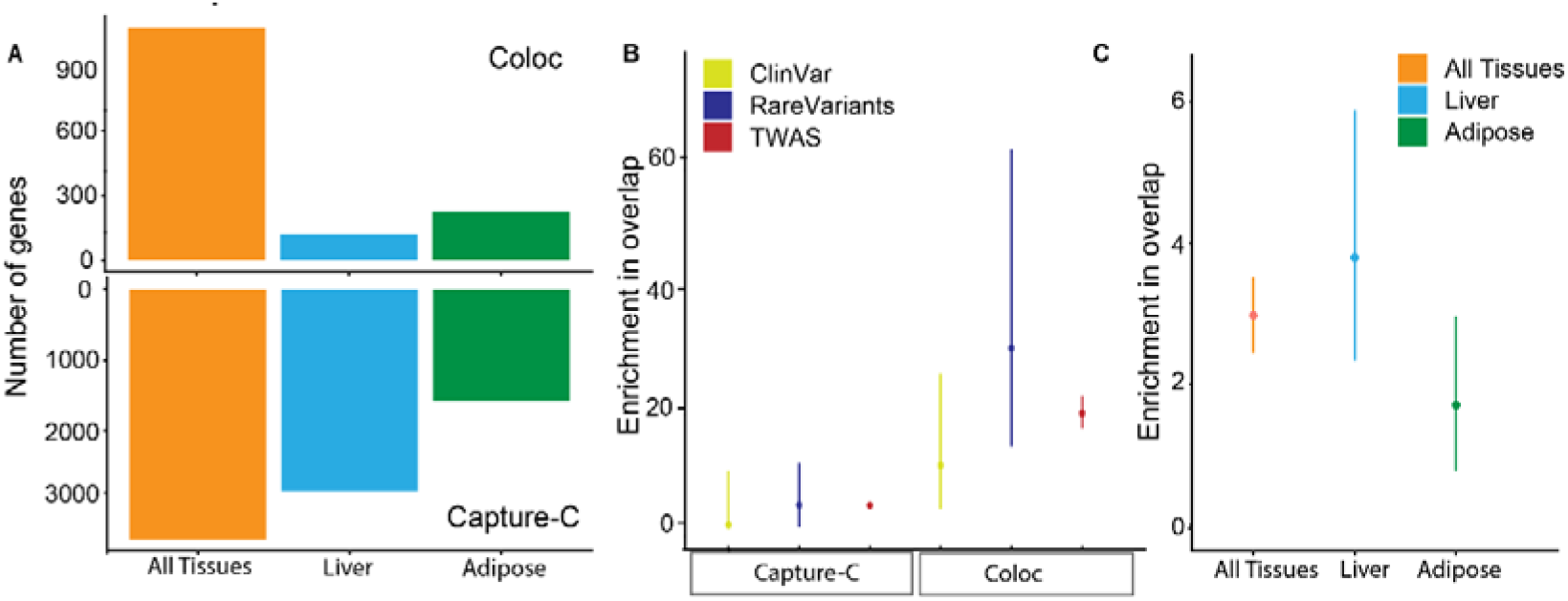
Overlap between eQTL colocalized genes and capture-C prioritized genes, and their enrichment in known lipid-associated genes. A. Numbers of genes identified by two approaches: eQTL colocalization (upper half) and promoter capture-c interactions (lower half) B. Overlap between our list of prioritized genes (left: capture-C prioritized genes; right: eQTL colocalized genes) with three sets of genes previously associated with lipid biology (ClinVar lipidemia-associated genes, genes implicated in rare burden of lipids, and genes from a lipid TWAS). C. Enrichment in overlap between eQTL colocalized genes and capture-C prioritized genes against what is expected by chance, assuming both gene sets are independent. Enrichment estimates and confidence intervals shown in Panels B and C were obtained using Fisher’s exact test.

To acquire additional functional insights into the 1,076 colocalized genes, we assessed their enrichments across existing biological and clinical gene sets. Colocalized genes showed enrichments in (a) 20 KEGG pathways ^30^ at FDR 5% (Table S2), including known lipid-related processes such as cholesterol metabolism, PPAR signaling, and bile secretion; (b) 33 Mendelian genes from ClinVar ^32^ associated with lipid-related ICD10 codes, (11 fold enrichment at P=2.08e-06, including *APOB, LPL*, and *APOE*; Figure 2B), suggesting the shared genetic basis of Mendelian and complex lipid phenotypes ^42^; (c) 35 genes with rare-variant burden for lipid phenotypes in a recent multi-ancestry analysis ^34^ (30-fold enrichment, P = 1.77e-16, including *APOB, LPL, LIPG* and *ANGPTL4*), confirming shared mechanisms of rare and common variation underlying lipid traits ^42,43^. Colocalized genes also showed enrichment with genes implicated in TWAS run on the same GWAS and eQTL summary statistics (20-fold enrichment, P<2.22e-308). These enrichment results demonstrate the biological relevance of candidate functional genes prioritized by our approach.

### Chromatin-chromatin interactions improve eQTL-based colocalization

Our eQTL-based colocalization analysis uses a linear sequence of DNA, and ignores physical interaction between non-adjacent DNA segments, another regulatory layer underlying complex human traits ^44^. To add this layer to our analysis, we generated Capture-C data from HepG2 liver carcinoma cells (denoted as HepG2.1) and hepatocyte-like cells (HLC) derived from differentiating iPSCs (the latter is described in ^22^), as well as publicly-available Capture -C datasets from HepG2 ^21,43^ (denoted as HepG2.2) and adipose tissue ^23^. We defined a GWAS-relevant interaction as any Capture-C interaction between any gene and a variant in the 95% credible set for a GWAS locus^45^. Credible set sizes ranged from 1 to 417 variants at the 1,750 examined loci, with a median size of 5 variants per credible set. In total, 1,079 GWAS loci had at least one variant in the credible set with a physical interaction with a gene promoter and 3,543 of 26,621 genes with promoter-interactions had promoters physically interacting with at least one GWAS credible set variant (Figure 2A; Table S3). Unlike eQTL-colocalized genes, these genes interacting with their credible sets showed limited enrichment in relevant KEGG pathways (Table S2) and lipid-related genes from ClinVar (Figure 2B), though we see 5-fold enrichment (compared to greater than 10-fold enrichment for eQTL-colocalized genes) in genes with rare-variant lipid associations (P =2.8e-05) and TWAS genes (P=2.5e-288).

Genes physically interacting with GWAS loci helped shortlist functional genes from eQTL colocalization despite their reduced enrichments in known gene sets. Of 1,079 credible sets with promoter interactions, 224 also colocalized with eQTLs for the same gene. At the gene level, 233 genes were implicated in both eQTL colocalization and Capture-C interactions (Figure 2C), representing an enrichment of 3-fold compared to random chance (P =3.11e-38). Among these loci with concordant eQTL colocalizations and Capture-C interactions, only 39% of them mapped to a single gene using eQTL data alone, whereas adding Capture-C information increased this fraction to 80%. These results showcase the potential value of combining eQTLs with physical chromatin interactions to prioritize functional genes at GWAS loci.

Since eQTLs are likely to reside in the same topologically associated domain (TADs) as the genes they regulate ^46^, we examined TAD structure from independent datasets at lipid GWAS loci with eQTL colocalizations. Of eQTL-GWAS colocalizations in which the sentinel variant resided within a liver TAD ^28^, the colocalized gene resided in the same liver TAD 84.8% of the time (P < 0.001 with 1000 permutations; Methods). When we restricted colocalizations to those supported by Capture-C data in any cell type, 91.2% fall in the same TAD. These results add to the existing evidence for TAD boundaries being regulatory insulators in the cell ^47^ and confirm our integration of chromatin interactions with eQTL colocalizations as an effective strategy to hone in on functional genes.

### Tissue-specific enrichment of GWAS signals differentiates lipid traits

Regulatory variants often affect complex traits in a tissue-specific manner ^6^, as shown in our eQTL colocalization analysis. Specifically, by computing the ratio of the number of colocalizations in a tissue to eQTL sample size in that tissue, we found that the liver was universally enriched for colocalized eGenes with respect to sample size across all lipid traits whereas adipose was selectively enriched in HDL and TG only (Figure S1). Motivated by these findings, we leveraged systematic approaches and additional data to identify relevant tissues and cell types for each lipid trait.

We implemented stratified LD score regression (S-LDSC), a polygenic approach not restricted to genome-wide significant variants, on tissue-specific transcriptomic and epigenomic annotations across 204 datasets from more than 170 tissues and cell types, to identify relevant tissues for each lipid trait (Methods). Consistent with previous studies and our eQTL-based analysis, liver-related tissues (Table S4) showed strong enrichments across all lipid traits (S-LDSC enrichment p-values ranging from .001 in TG to .0001 in TC), for both expression (Figure 3A) and chromatin annotations (Figure 3B). This result was confirmed by analysis using two other approaches: DEPICT ^48^ (Figure S2) and RSS-NET ^49^ (Table S5). To assess the robustness of our S-LDSC results based on multi-population GWAS, we applied S-LDSC to population-specific GWAS in European and East Asian ancestry participants together with population-specific LD scores (Methods) and obtained similar results (Table S6).

**Figure 3:**
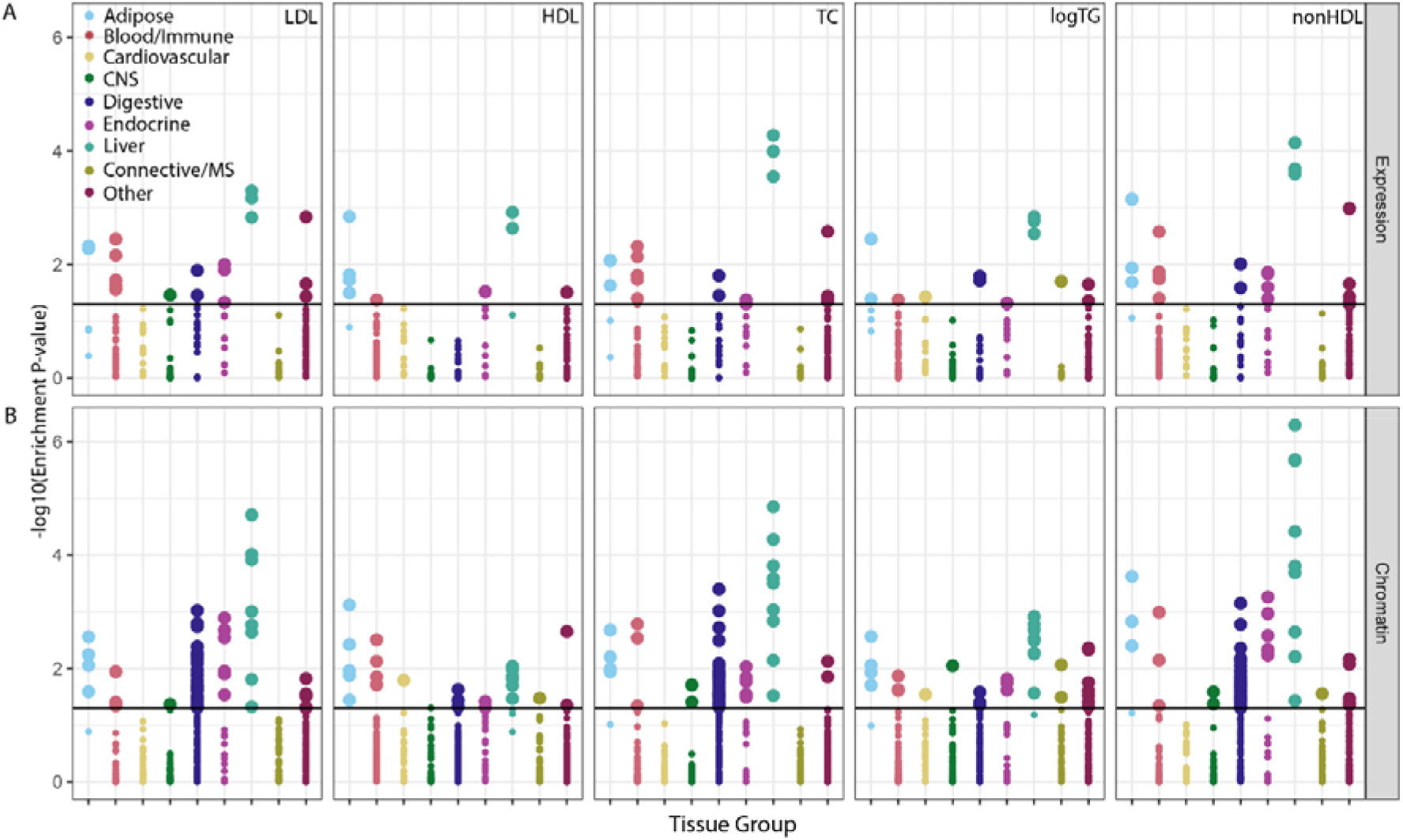
Tissue relevance based on lipid GWAS and functional annotations. Partitioning heritability of GWAS summary statistics for five lipid traits on gene expression (A) and chromatin mark (B) annotations across tissues. Each plotted point represents a tested dataset for enrichment of heritability, with larger dots representing datasets with P-value < 0.05; multiple annotation datasets are tested for the same tissue group. Each color represents a tissue group, and the y-axis represents -log10 P-value of enrichment of heritability.

The S-LDSC results also highlighted tissues selectively enriched in certain lipid traits as shown in the eQTL-based analysis. The most enriched category for HDL using chromatin annotation is ‘Adipose H3K4me3’ (P-value 7.6e-04); for TG, enrichment in liver-related tissues (P-value 1.2e-03) is similar to enrichment in adipose (P-value 2.7e-03). For LDL, TC, and non-HDL, enrichment P-values for the liver were much more significant than for all other tissues including adipose (Figure 3B). We observed the same pattern in S-LDSC results based on gene expression (Figure 3A). This finding is consistent with the known influence of adipose on plasma HDL levels ^50^, and the role of adipose as TG deposits ^51^. These results were corroborated by eQTL colocalizations stratified by phenotype (Figure S1) and DEPICT analysis on gene expression ^48^ (Figure S2). Together, these results confirm the liver as the tissue of action for all five lipid traits, and highlight the additional role of adipose in HDL and TG only.

Given the importance of the liver and adipose in modulating lipid levels, we further identified the relevant cell types within these tissues. Using existing single-cell data from adipose and liver, we performed gene-set enrichment analysis ^52^ to identify cell-type clusters enriched for genes colocalized with any lipid trait. Out of 11 identified cell types in 20 clusters in the liver, only hepatocytes were enriched at FDR-adjusted P < 0.05 (Figure S3), consistent with previous results^21^. In adipose, only adipocyte clusters and macrophage-monocyte clusters showed suggestive enrichment (nominal P < 0.05) in colocalized genes (Figure S4). Of note, the enrichment in adipocytes was significant when we restricted this analysis to genes that were colocalized only with HDL and TG (FDR-corrected P < 0.05), consistent with the selective enrichments of adipose in HDL and TG (but not the other lipid traits) from our S-LDSC analysis. Evaluations at cellular resolution are required to understand the cell-type specific mechanisms underlying lipid GWAS loci, but our results could form a useful basis for future studies.

### Overlapping GWAS signals with binding sites highlights lipid-relevant TFs

TFs have been implicated as a key mediator of linking genetic variation to complex traits ^53^. To understand lipid GWAS in the context of TF activity, we assessed enrichment of genome-wide significant variants at TF binding sites using GREGOR ^37^ and performed polygenic enrichment analysis of TF binding sites using S-LDSC.

Using ChIP-Seq data from 161 TFs across 91 cell types from the ENCODE project ^7^, 70.7% of lipid credible sets overlapped with at least one TF binding site. Using GREGOR ^37^, we identified 137 TFs whose binding sites were significantly enriched in GWAS lead SNPs for at least one lipid phenotype (enrichment > 2; FDR adjusted P-value < 0.05; Figure S5; Table S7). Among these 137 enriched TFs, 69 of them (50%) showed significant enrichments across all five lipid phenotypes, suggesting a potential core regulatory circuit shared by all lipid traits (Figure S5). The TF with the strongest enrichment in all phenotypes was ESRRA (estrogen-related receptor alpha), a nuclear receptor active in metabolic tissues ^54^; ESRRA has been implicated in adipogenesis and lipid metabolism, and ESRRA-null mice display an increase in fat mass and obesity ^54^.

The GREGOR analysis also highlighted 68 TFs significantly enriched in specific subsets of (but not all five) lipid phenotypes (Figure S8). For example, we found 4 TFs (FOXM1, PBX3, ZKSCAN1, ZEB1) enriched in HDL and TG only, 4 TFs (EZH2, NFE2, NFATC1, KDM5A) enriched in HDL only and 11 TFs (FOSL1, IRF3, JUN, MEF2C, NANOG, PRDM1, RUNX3, SIRT6, SMC3, STAT3, ZNF217) enriched in TG only. Of these TFs, the central role of ZEB1 in adiposity ^55^ and fat cell differentiation has been demonstrated ^56^. Taken together, these TF-centric findings corroborate the selective enrichments of adipose in HDL and TG (but not the other lipid traits) identified in our previous tissue prioritization analyses.

Similar to tissue prioritization, we also performed polygenic enrichment analysis of TF binding sites using S-LDSC (Table S8), which differed from GREGOR analysis by looking at not only the genome-wide significant associations but also the polygenic signal irrespective of GWAS P-values. On the same 161 ENCODE TFs, this polygenic analysis identified 25 TFs whose binding sites were significantly enriched in heritability (nominal P < 0.05) for at least one lipid phenotype (Figure S6); reassuringly, 24 of 25 TFs were also significant in GREGOR analysis. Among these enriched TFs, eight (34%) were significantly enriched in all five lipid traits (CEBPB, CEBPD, FOXA2, HDAC2, HNF4G, NFYA, RXRA, SP1; P < 0.05). Of those TFs significant in both analyses, RXRA (retinoid X receptor alpha) is also encoded by a colocalized gene (*RXRA*) near a GWAS hit (chr9:137,268,682). RXRA is a ligand-activated transcription factor that forms heterodimers with other receptors (including PPARG) and is involved in lipid metabolism ^57^ and homeostasis. Moreover, 145 GWAS loci (Table S9) overlap RXRA binding peaks, suggesting that the GWAS variants might affect lipids (partially) through affecting the binding activity of RXRA. While the *RXRA*-associated variant has been previously implicated as a GWAS locus ^58^, our study demonstrates its role in lipid biology through its regulatory influence on other lipid-associated genes.

### Multi-layer functional integration reveals regulatory mechanisms at GWAS loci

Motivated by our finding that integrating chromatin interaction improved eQTL colocalizations, we further brought together multiple lines of functional evidence at each GWAS locus for mechanistic inference. We started with the list of genes with evidence for both eQTL colocalization in the liver or adipose and credible set physical interactions. We next annotated each variant in the 95% credible set with various indicators of regulatory function, including its open chromatin status in liver or adipose-related cell types, its proximity to a promoter or an enhancer, and its RegulomeDB regulation probability ^59^ (see Table S10 for the complete list of annotations used). To account for complexities of regulatory mechanisms and limitations of functional datasets, we combined evidence across these datasets to prioritize variants at GWAS loci (Figure 4A). Specifically, we prioritized variants with at least three independent lines of functional evidence (chromatin openness, physically interaction with target genes, and promoter/enhancer status in liver or adipose), with at least two being in the same tissue with colocalization with the target gene, and with a RegulomeDB score > 0.5. Applying this simple procedure to lipid GWAS we identified 13 candidate loci, each with the strongest multi-layer evidence pointing to a single functional variant (Table 1). Below we describe two examples to highlight key features of this multi-layer integration framework.

**Figure 4.**
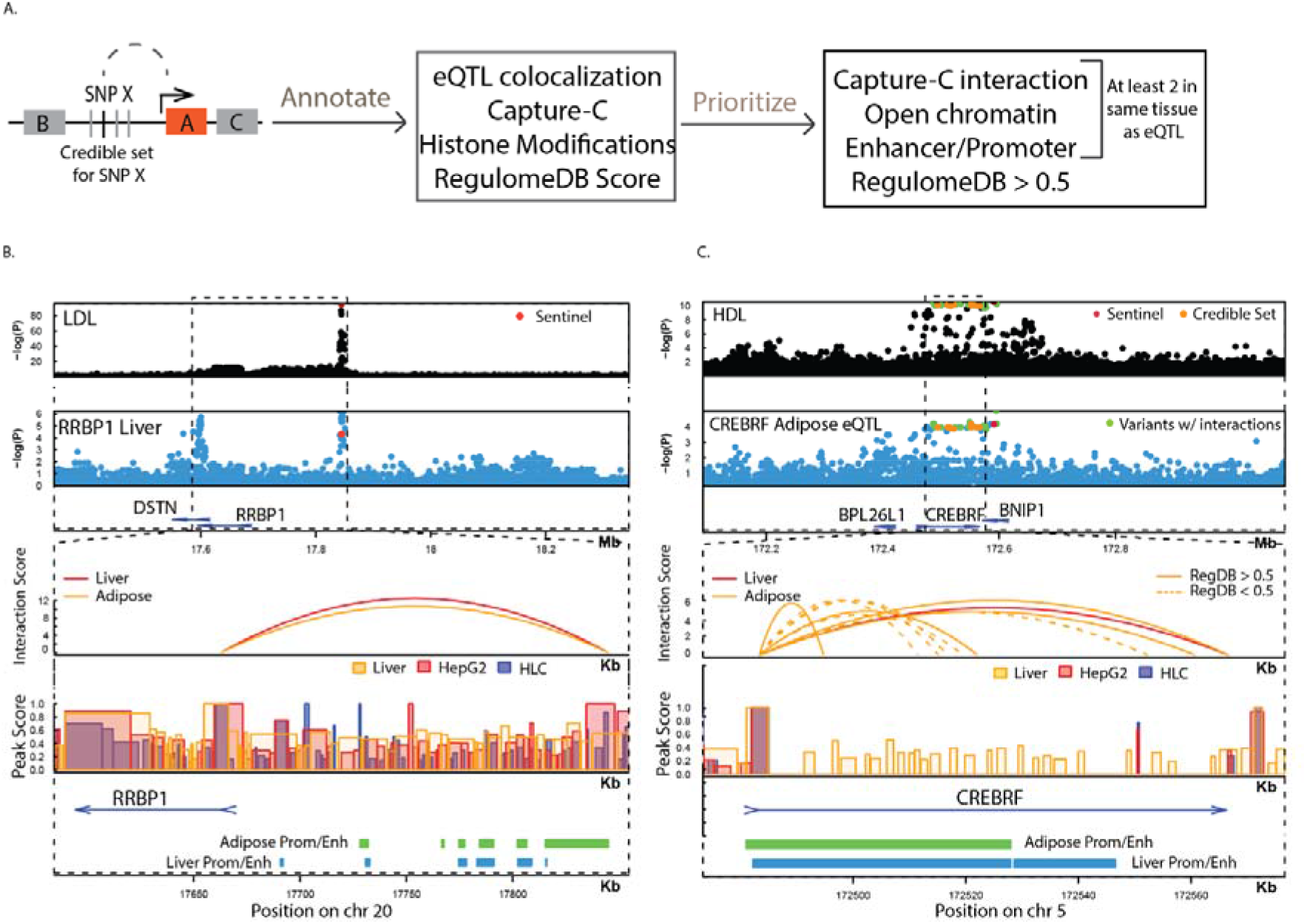
An easy-to-implement multi-layer framework to prioritize functional variants at GWAS loci. A. Variant annotation and prioritization scheme at each credible set. B. Evidence for gene RRBP1 from functional genomics data. The LDL GWAS locus at this region is an eQTL for gene RRBP1 in the liver (second row). Variants in the credible set of this locus interact with the gene promoter in both adipose and HepG2 Capture-C data. The interacting variant is also in an open chromatin peak in three liver-related cell types. C. Multiple sources of functional genomics data support CREBRF as a gene contributing to HDL levels. The HDL GWAS locus at this region is an eQTL for gene CREBRF in adipose (second row). Variants in the credible set at this locus interact with the CREBRF promoter in adipose. The interacting variant is also in open chromatin in liver-related cell types.

**Table 1.** Thirteen prioritized loci with highest confidence of a single functional variant in the credible set. The ‘sentinel’ column represents the lead variant at the locus. ‘Prioritized var’ represents the prioritized variant in the credible set. Columns 5-8 represent overlap of the functional variant with open chromatin (‘Open’), capture-C (‘CapC’) interactions with the candidate gene, enhancer and promoter marks from Roadmap in liver (‘Liver’), adipose (‘Ad’), both or none of these datasets. The ‘RegDB’ column represents the RegulomeDB score of the prioritized variant.

| *Gene Name* | *Tissue* | *Sentinel* | *Prioritized Var* | *Open* | *CapC* | *Enhancer* | *Promoter* | *RegDB* |
| --- | --- | --- | --- | --- | --- | --- | --- | --- |
| *CEP68* | Adipose | 2:65284231 | 65279414 | Liver | Liver | None | Ad | 0.5896 |
| *TIPARP* | Adipose | 3:156797941 | 156795408 | Both | Both | Ad | Liver | 0.705 |
| *CREBRF* | Adipose | 5:172591337 | 172566698 | Liver | Ad | None | Both | 0.9124 |
| *PALM2* | Adipose | 9:112556911 | 112556911 | Both | Ad | Both | None | 0.6091 |
| *MEGF9* | Adipose | 9:123481206 | 123421556 | Liver | Ad | None | Liver | 0.9933 |
| *GBF1* | Liver | 10:104142294 | 104107191 | Ad | Ad | Both | Ad | 0.705 |
| *MICAL2* | Liver | 11:12071855 | 12221016 | Liver | Liver | Liver | Ad | 0.6018 |
| *ACP2* | Liver | 11:47278917 | 47276350 | Ad | Liver | Liver | Ad | 0.6091 |
| *PTPRJ* | Adipose | 11:48021778 | 48011180 | Liver | Ad | Liver | Ad | 0.8797 |
| *NFATC2IP* | Adipose | 16:28899411 | 28883327 | Liver | Liver | None | Both | 0.6091 |
| *HELZ* | Liver | 17:65109591 | 65156919 | Liver | Liver | Both | Ad | 0.60906 |
| *FAM210A* | Liver | 18:13725674 | 13725674 | Liver | Liver | Both | Ad | 0.7571 |
| *RRBP1* | Liver | 20:17844684 | 17844684 | Both | Ad | Both | Ad | 0.6091 |

*RRBP1* (ribosomal binding protein 1) could be identified from eQTL colocalization alone, but our multi-layer integration approach strengthened the conclusion via convergent evidence from various sources (Figure 4B). The *RRBP1* eQTL signals in the liver colocalize with LDL, TC, and nonHDL GWAS signals. The ‘T’ allele of the lead variant (chr20:17,844,684, hg19) decreases *RRBP1* expression levels and increases LDL, TC, and nonHDL levels. This lead variant is in open chromatin in HLC, and physically interacts with the *RRBP1* promoter (250kb away) in adipose and HepG2. All these data consistently point to *RRBP1* as the functional gene underlying this locus. RRBP1 specifically tethers the endoplasmic reticulum to the mitochondria in the liver--an interaction that is enriched in hepatocytes--and regulates very low density lipoprotein (vLDL) levels ^60^. Rare variants in *RRBP1* are associated with LDL in humans ^61^ and silencing *RRBP1* in liver affects lipid homeostasis in mice ^60^.

*CREBRF* (CREB3 regulatory factor) demonstrates the power of our multi-layer integration framework in prioritizing functional variants (Figure 4C). The eQTL signals of *CREBRF* colocalized with a GWAS locus for HDL with 30 candidate variants. In contrast, our multi-layer approach identified a single candidate variant (chr5:172,566,698) at this locus that physically interacts with the *CREBRF* promoter in adipose, was predicted to be a regulatory element (RegulomeDB score=0.91). Consistent with the index variant (chr5:172,591,337), the allele ‘A’ at this functional variant increased HDL levels and increased *CREBRF* expression in adipose. Missense variants in *CREBRF* have been linked to body mass index, and the gene has been linked to obesity risk in Samoans ^62^.

Finally, to compare the power of functional fine-mapping with trans-ancestry fine-mapping, we applied our prioritization rule to credible sets derived from European-only meta-analysis. The 111 variants prioritized by our rule described above (including multiple variants in the same credible set) were all found in the multi-ancestry credible sets, representing a 3.7 fold enrichment (P < 1e-04 derived from 10000 permutations randomly sampling variants from the European-only credible sets). This convergence of complementary approaches to the same smaller set of variants highlights the power of multi-ancestry datasets as an approach to narrow in on functional variants.

## Discussion

Here we integrate the largest multi-population lipid GWAS to date with a wide array of functional genomic resources to understand how noncoding genetic variation affects lipids through gene regulation. Specifically, we identify 1,076 genes whose eQTL signals colocalize with lipid GWAS signals and demonstrate how physical chromatin interaction can improve standard eQTL-based colocalization. We assess tissue-specific enrichments of lipid GWAS signals and demonstrate the selective importance of adipose in HDL and triglyceride biology. We examine binding site enrichments of 161 TFs in lipid GWAS and expand our understanding of lipid GWAS loci (e.g., *RXRA*) in the context of TF activity. Finally, we build a simple and interpretable prioritization framework that automatically combines multiple lines of evidence from orthogonal datasets, pinpointing a single functional variant at each of 13 lipid-associated loci (e.g., *RRBP1* and *CREBRF*). While there are studies that interpret lipid GWAS associations ^21,63,64^, the size of our multi-population GWAS and multi-layer functional integration represent a comprehensive effort and an important step forward in this direction.

Our multi-layer analysis has two key strengths. First, despite a large array of functional genomic resources being embedded, our analysis produces results with high consistency. For example, the selective enrichment of adipose in HDL and TG identified by S-LDSC is confirmed by our eQTL-based colocalization and TF binding site overlap. Another example of consistency is the multi-layer prioritization of *RRBP1*, which can be identified from eQTL-based colocalization alone and it is further validated by chromatin openness and interaction. Such convergent evidence from various sources improves the confidence of our findings. Second, our analysis highlights that combining multiple layers of regulatory information can improve sensitivity to prioritize functional genes and variants. For example, we refined eQTL colocalized genes (1,076) to a smaller set of functional genes (233) through integration with promoter Capture-C data. Another example of sensitivity is *CREBRF*, where eQTL-based colocalization implicates 30 candidate variants and adding other regulatory layers points to a single functional variant. Moving forward, we expect these two features will serve as useful guidelines for future integrative genomic analyses of other traits.

Our results rely on the breadth and accuracy of functional genomic datasets used in our analyses. First, unlike our lipid GWAS, current functional datasets ^65^ are limited both in sample size and ancestral diversity, which can affect discovery and replication of regulatory mechanisms in diverse populations. Second, some functional datasets are generated at limited resolution. For example, our colocalizations are based on eQTLs from bulk tissue RNA-seq ^8^, which may miss detailed cell types and biological processes in which lipid-associated SNPs regulate gene expression ^66^. Third, some functional datasets are not available across the full spectrum of human tissues and cell types. For example, our chromatin-chromatin interaction analysis only examines a few cell types in two known lipid-related tissues, producing results that may be biased towards known lipid biology. As more comprehensive and accurate functional genomic resources are becoming publicly available in diverse cellular contexts and ancestry groups, the resolution and power of integrative analyses like ours will be markedly increased.

Other limitations of this study stem from computational methods embedded in our framework. First, the colocalization approach ‘coloc’ assumes one causal variant per locus, whereas recent studies suggest extensive allelic heterogeneity ^67^ consistent with a model of a milieu of related transcription factors binding within a single locus. Accounting for allelic heterogeneity in summary statistics-based colocalization typically requires modelling multiple correlated SNPs through LD matrix ^68^, which is computationally intensive in large-scale analyses derived from many cohorts with diverse ancestries, like the multi-population GWAS examined here. Second, due to restricted access to individual genotypes of 201 cohorts, we cannot produce multi-population LD scores within GLGC but have to use European-based LD scores in all S-LDSC analyses. This approach, though less rigorous in principle, provides robust results in practice (as confirmed by our ancestry-specific analysis), largely because 79% of cohorts in GLGC are of European descent ^12^. That said, we caution that the same approach might fall short in ancestrally diverse studies with few European individuals ^69^. Third, our multi-layer variant prioritization framework is built on a series of simple rules that are easy to implement on large datasets. This approach could possibly be formalized as statistical models (e.g., priors in Bayesian methods ^49^), but certainly simplify computation and improve scalability of our framework. Despite the technical limitations, our approach here can serve as a useful benchmark for future development of methods with improved statistical rigor and computation efficiency.

In summary, mapping noncoding genetic variation of complex traits to biological functions can benefit greatly from thorough integration of multiple layers of functional genomics, as demonstrated in the present study. Although tested on lipids only, our integrative framework is straightforward to implement more broadly on many other phenotypes, yielding functional insights of heritable traits and diseases in humans.

## Description of Supplemental Data

Supplemental data include seven figures and ten tables, and study-specific acknowledgements.

## Acknowledgments

GMP, PN and CW are supported by NHLBI R01HL127564. GMP and PN are supported by R01HL142711. AG acknowledge support from the Wellcome Trust (201543/B/16/Z), European Union Seventh Framework Programme FP7/2007-2013 under grant agreement no. HEALTH-F2-2013-601456 (CVGenes@Target) & the TriPartite Immunometabolism Consortium [TrIC]-Novo Nordisk Foundation’s Grant number NNF15CC0018486. JMM is supported by American Diabetes Association Innovative and Clinical Translational Award 1-19-ICTS-068. SR was supported by the Academy of Finland Center of Excellence in Complex Disease Genetics (Grant No 312062), the Finnish Foundation for Cardiovascular Research, the Sigrid Juselius Foundation and University of Helsinki HiLIFE Fellow and Grand Challenge grants. EW was supported by the Finnish innovation fund Sitra (EW) and Finska Läkaresällskapet. CNS was supported by American Heart Association Postdoctoral Fellowships 15POST24470131 and 17POST33650016. Charles N Rotimi is supported by Z01HG200362. Zhe Wang, Michael Presuss, and Ruth JF Loos are supported by R01HL142302. NJT is a Wellcome Trust Investigator (202802/Z/16/Z), is the PI of the Avon Longitudinal Study of Parents and Children (MRC & WT 217065/Z/19/Z), is supported by the University of Bristol NIHR Biomedical Research Centre (BRC-1215-2001), the MRC Integrative Epidemiology Unit (MC_UU_00011) and works within the CRUK Integrative Cancer Epidemiology Programme (C18281/A19169). Ruth E Mitchell is a member of the MRC Integrative Epidemiology Unit at the University of Bristol funded by the MRC (MC_UU_00011/1). Simon Haworth is supported by the UK National Institute for Health Research Academic Clinical Fellowship. Paul S. de Vries was supported by American Heart Association grant number 18CDA34110116. Julia Ramierz acknowledges support by the People Programme of the European Union’s Seventh Framework Programme grant n° 608765 and Marie Sklodowska-Curie grant n° 786833. Maria Sabater-Lleal is supported by a Miguel Servet contract from the ISCIII Spanish Health Institute (CP17/00142) and co-financed by the European Social Fund. Jian Yang is funded by the Westlake Education Foundation. Olga Giannakopoulou has received funding from the British Heart Foundation (BHF) (FS/14/66/3129). Dr. Sander W. van der Laan is funded through grants from the Netherlands CardioVascular Research Initiative of the Netherlands Heart Foundation (CVON 2011/B019 and CVON 2017-20: Generating the best evidence-based pharmaceutical targets for atherosclerosis [GENIUS I&II]). We are thankful for the support of the FP7 EU project CVgenes@target (HEALTH-F2-2013-601456), ERA-CVD program ‘druggable-MI-targets’ (grant number: 01KL1802) and the Leducq Fondation ‘PlaqOmics’. Study-specific acknowledgements are available in the **Supplementary Material**.

## Web Resources

GLGC 2021 summary statistics: http://csg.sph.umich.edu/willer/public/glgc-lipids2021/

GTEx v8 summary statistics: https://www.gtexportal.org/home/datasets

coloc: https://cran.r-project.org/web/packages/coloc

liftOver: https://genome.ucsc.edu/cgi-bin/hgLiftOver

HiCUP: https://www.bioinformatics.babraham.ac.uk/projects/hicup/

CHiCAGO: https://www.bioconductor.org/packages/release/bioc/html/Chicago.html

GenomicRanges: https://bioconductor.org/packages/release/bioc/html/GenomicRanges.html

bedtools: https://bedtools.readthedocs.io/en/latest/

ClusterProfiler: https://guangchuangyu.github.io/clusterProfiler

biomaRt: https://bioconductor.org/packages/release/bioc/html/biomaRt.html

ClinVar: https://www.ncbi.nlm.nih.gov/clinvar/

S-PrediXcan: https://github.com/hakyimlab/MetaXcan

LDSC software: https://github.com/bulik/ldsc

LD scores and related annotations: https://data.broadinstitute.org/alkesgroup/LDSCORE/

DEPICT: https://data.broadinstitute.org/mpg/depict

RSS-NET: https://github.com/SUwonglab/rss-net

Adipose single cell data: https://singlecell.broadinstitute.org/single_cell/study/SCP133/human-adipose-svf-single-cell

fgsea: http://bioconductor.org/packages/release/bioc/html/fgsea.html

GREGOR: https://genome.sph.umich.edu/wiki/GREGOR

RegulomeDB: https://regulomedb.org/regulome-search/

## Data and Code Availability

HLC Capture-C data is available at https://www.ncbi.nlm.nih.gov/geo/query/acc.cgi?acc=GSE189026

