## Supplementary Figures for "A multi-layer functional genomic analysis to understand noncoding genetic variation in lipids"

*Figure S1.* Number of colocalizations per tissue (Y axis) is directly proportional to the eQTL sample size (X axis) for that tissue. The red dot is the liver, the blue dot is adipose subcutaneous.

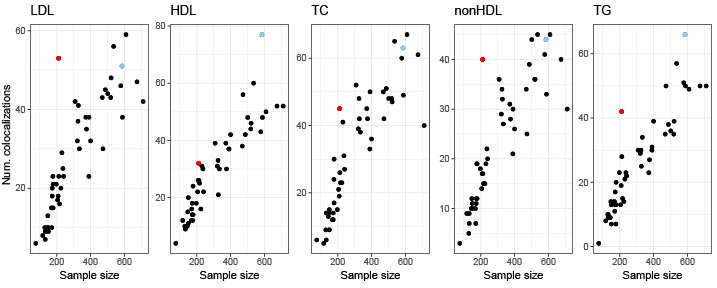

*Figure S2*. DEPICT analysis on GWAS summary statistics shows different tissues enriched for each lipid phenotype. Adipose-related tissues are particularly enriched in HDL and logTG compared to the liver, which is enriched for all phenotypes.

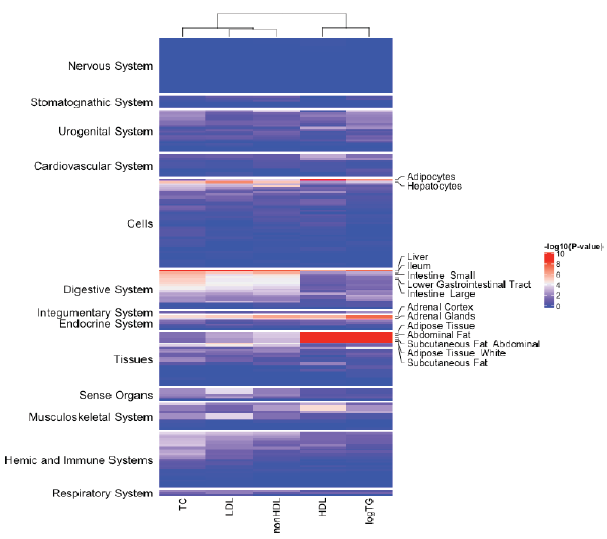

*Figure S3*. Hierarchical clustering of normalized gene expression for colocalized genes in hepatic cell types (10 cell types organized into 20 single-cell clusters). Hepatocytes (labeled ‘Hep’) are most strongly enriched in gene expression (red shows stronger expression and blue shows weaker). Cell types are labeled as in MacPharland et al, 2018.

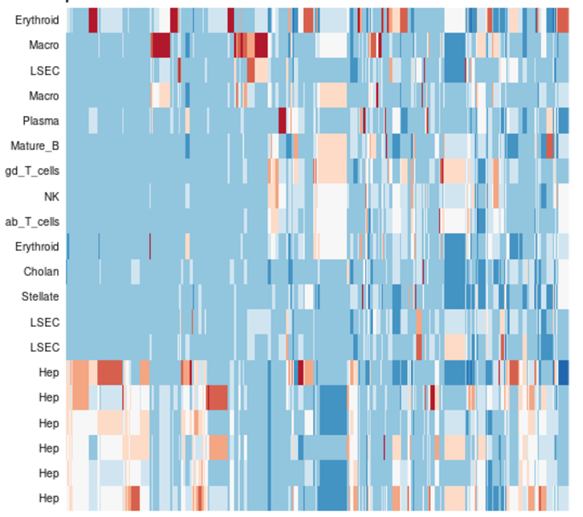

*Figure S4*. Heatmap of normalized gene expression for colocalized genes in adipose cell types (10 cell types organized into 20 single-cell clusters). Cell type names are (monocytes; adipocytes; dendritic; keratinocytes; Bcells; mast cells; T-cells; NK cells; vascular cells; Lymphatic endothelial cells; preadipocytes)

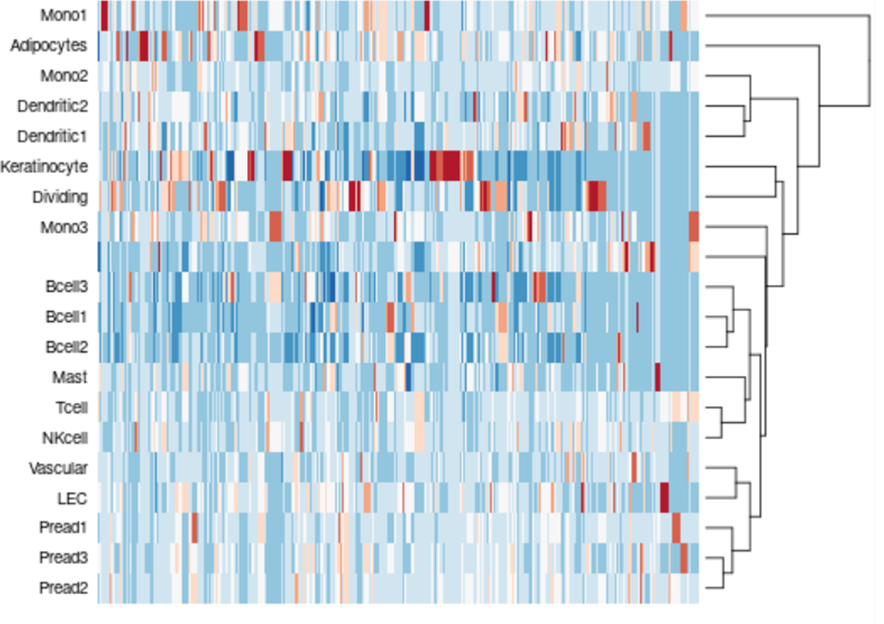

*Figure S5.* Enrichments in TF binding in significant associations for each phenotype. We see a strong correlation in enriched TFs across lipid phenotypes, and highlight TFs showing the strongest enrichments

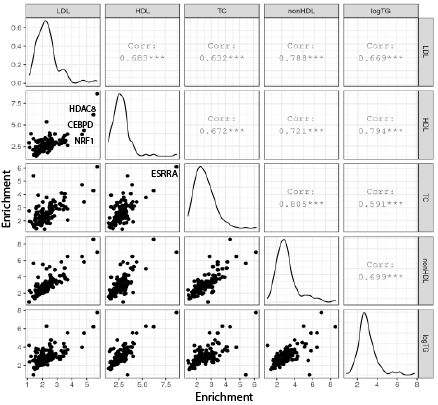

*Figure S6*: Number of TFs showing enrichment in GWAS signal for each phenotype using GREGOR (top), and s-LDSC (bottom)

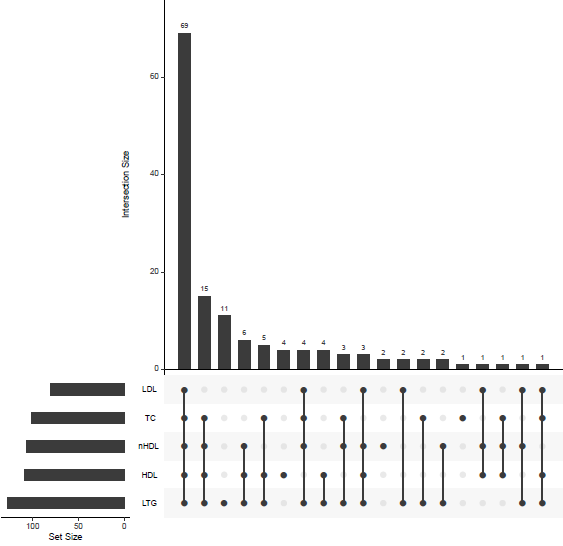

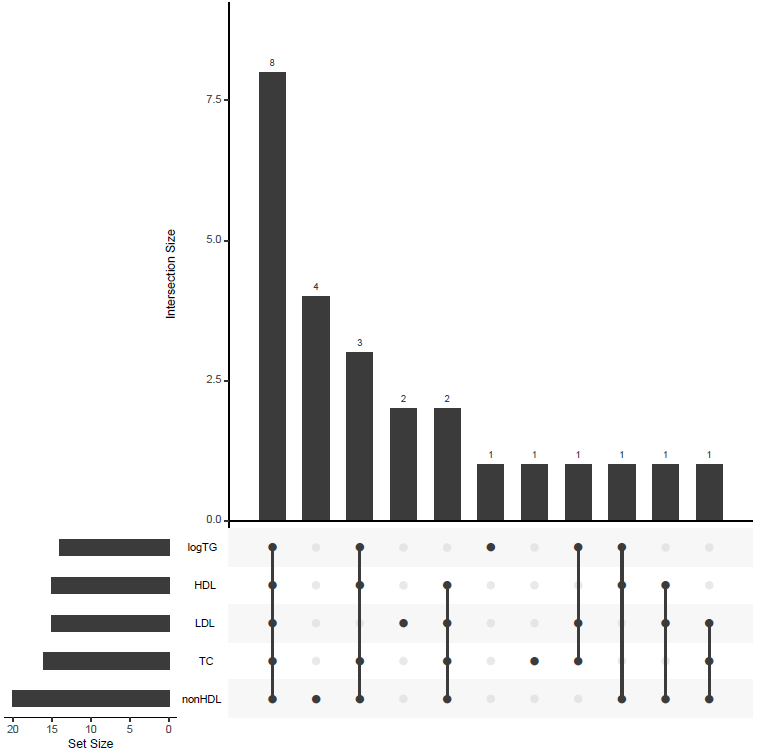

**Supplementary Acknowledgements by Cohort:**

Africa America Diabetes Mellitus

The authors would like to gratefully acknowledge the AADM collaborators, including Olufemi Fasanmade, Thomas Johnson, Johnnie Oli, Godfrey Okafor, Benjamin A. Eghan Jr., Kofi Agyenim-Boateng, Clement Adebamowo, Albert Amoah, Joseph Acheampong, and Duncan Ngare (deceased). This project was supported by the Intramural Research Program of the National Human Genome Research Institute of the National Institutes of Health (NIH) through the Center for Research on Genomics and Global Health (CRGGH). The CRGGH is also supported by the National Institute of Diabetes and Digestive and Kidney Diseases and the Office of the Director at the NIH (Z01HG200362). Support for participant recruitment and initial genetic studies of the AADM study was provided by NIH grant No. 3T37TW00041-03S2 from the Office of Research on Minority Health. This work utilized the computational resources of the NIH HPC Biowulf cluster. (<http://hpc.nih.gov>)

Age-related diseases: Understanding Genetic and non-genetic influences - a study at the University of Regensburg

Cohort recruiting and management was funded by the Federal Ministry of Education and Research (BMBF-01ER1206, BMBF-01ER1507, to Iris M. Heid) and by the German Research Foundation (DFG HE 3690/7-1, to Iris M. Heid). Genome-wide genotyping and lipid concentrations phenotyping was funded by the University of Regensburg for the Department of Genetic Epidemiology. The computational work supervised by Iris M. Heid was funded by the German Research Foundation (DFG) – Project-ID 387509280 – SFB 1350 (to Iris M. Heid).

Anglo-Scandinavian Cardiac Outcomes Trial

This work was funded by the National Institutes for Health Research (NIHR) as part of the portfolio of translational research of the NIHR Barts Biomedical Research Unit and the NIHR Biomedical Research Centre at Imperial College, the International Centre for Circulatory Health Charity and the Medical Research Council through G952010. We thank all ASCOT trial participants, physicians, nurses, and practices in the participating countries for their important contribution to the study. The study was investigator-led and was conducted, analyzed, and reported independently of Pfizer who funded the trial. JR acknowledges support by the People Programme of the European Union’s Seventh Framework Programme grant n° 608765 and Marie Sklodowska-Curie grant n° 786833.

Atherosclerosis Risk in Communities Study

The Atherosclerosis Risk in Communities study has been funded in whole or in part with Federal funds from the National Heart, Lung, and Blood Institute, National Institutes of Health, Department of Health and Human Services (contract numbers HHSN268201700001I, HHSN268201700002I, HHSN268201700003I, HHSN268201700004I and HHSN268201700005I), R01HL087641, R01HL059367 and R01HL086694; National Human Genome Research Institute contract U01HG004402; and National Institutes of Health contract HHSN268200625226C. The authors thank the staff and participants of the ARIC study for their important contributions. Infrastructure was partly supported by Grant Number UL1RR025005, a component of the National Institutes of Health and NIH Roadmap for Medical Research. Paul S. de Vries was supported by American Heart Association grant number 18CDA34110116.

Austrian Stroke Prevention Study/Austrian Stroke Prevention Family Study

The authors thank the staff and the participants for their valuable contributions. We thank Birgit Reinhart for her long-term administrative commitment, Elfi Hofer for the technical assistance at creating the DNA bank, Ing. Johann Semmler and Anita Harb for DNA sequencing and DNA analyses by TaqMan assays and Irmgard Poelzl for supervising the quality management processes after ISO9001 at the biobanking and DNA analyses.

The Medical University of Graz and the Steiermärkische Krankenanstaltengesellschaft support the databank of the ASPS/ASPS-Fam. The research reported in this article was funded by the Austrian Science Fund (FWF) grant numbers PI904, P20545-P05 and P13180 and supported by the Austrian National Bank Anniversary Fund, P15435 and the Austrian Ministry of Science under the aegis of the EU Joint Programme-Neurodegenerative Disease Research (JPND)-www.jpnd.eu.

Avon Longitudinal Study of Parents and Children

NJT is a Wellcome Trust Investigator (202802/Z/16/Z), is the PI of the Avon Longitudinal Study of Parents and Children (MRC & WT 217065/Z/19/Z), is supported by the University of Bristol NIHR Biomedical Research Centre (BRC-1215-2001), the MRC Integrative Epidemiology Unit (MC_UU_00011) and works within the CRUK Integrative Cancer Epidemiology Programme (C18281/A19169). SH is supported by the UK National Institute for Health Research Academic Clinical Fellowship. REM is a member of programme 1 at the MRC Integrative Epidemiology Unit at the University of Bristol funded by the MRC (MC_UU_00011/1)

the Beijing Atherosclerosis Study

This work was supported by the CAMS Innovation Fund for Medical Sciences (grants 2019-I2M-2-003, 2016-I2M-1-009, 2017-I2M-1-004, and 2016-I2M-1-011).

BioBank Japan

We acknowledge the staff of the BBJ for their outstanding assistance. This research was supported by the Tailor-Made Medical Treatment Program (BBJ) of the Ministry of Education, Culture, Sports, Science and Technology (MEXT), the Japan Agency for Medical Research and Development (AMED) under grant number JP17km0305002.

British 1958 birth cohort

We acknowledge use of phenotype and genotype data from the British 1958 Birth Cohort DNA collection, funded by the Medical Research Council grant G0000934 and the Wellcome Trust grant 068545/Z/02. Genotyping for the B58C-WTCCC subset was funded by the Wellcome Trust grant 076113/B/04/Z. The B58C-T1DGC genotyping utilized resources provided by the Type 1 Diabetes Genetics Consortium, a collaborative clinical study sponsored by the National Institute of Diabetes and Digestive and Kidney Diseases (NIDDK), National Institute of Allergy and Infectious Diseases (NIAID), National Human Genome Research Institute (NHGRI), National Institute of Child Health and Human Development (NICHD), and Juvenile Diabetes Research Foundation International (JDRF) and supported by U01 DK062418. B58C-T1DGC GWAS data were deposited by the Diabetes and Inflammation Laboratory, Cambridge Institute for Medical Research (CIMR), University of Cambridge, which is funded by Juvenile Diabetes Research Foundation International, the Wellcome Trust and the National Institute for Health Research Cambridge Biomedical Research Centre; the CIMR is in receipt of a Wellcome Trust

Strategic Award (079895). The B58C-GABRIEL genotyping was supported by a contract from the European Commission Framework Programme 6 (018996) and grants from the French Ministry of Research.

British Genetics of Hypertension Trial

This work was funded by the Medical Research Council of Great Britain (grant number: G9521010D). The BRIGHT study is extremely grateful to all the patients who participated in the study and the BRIGHT nursing team. This work forms part of the research themes contributing to the translational research portfolio for the NIHR Barts Cardiovascular Biomedical Research Unit. The funders had no role in study design, data collection and analysis.

CAGE-KING

The KING Study was supported in part by Grants-in-Aid from MEXT (nos. 24390169, 16H05250, 25293144, 15K19242, 16H06277, 19K19434, 20K10514, 21H03206) as well as by a grant from the Funding Program for Next-Generation World-Leading Researchers (NEXT Program, no. LS056)

CARDIOGENICS

The main sponsor of Cardiogenics was the EU (LSHM-CT-2006-037593). This work was supported by the British Heart Foundation (BHF) grant RG/14/5/30893 (P.D.) and forms part of the research themes contributing to the translational research portfolios of the Barts Biomedical Research Centre funded by the UK National Institute for Health Research (NIHR)

Cardiovascular Health Study

The CHARGE cohorts were supported in part by R01HL105756. This CHS research was supported by contract 75N92021D00006**,** NHLBI contracts HHSN268201200036C, HHSN268200800007C, HHSN268201800001C, N01HC55222, N01HC85079, N01HC85080, N01HC85081, N01HC85082, N01HC85083, N01HC85086; and NHLBI grants U01HL080295, R01HL087652, R01HL105756, R01HL103612, R01HL120393, and U01HL130114 with additional contribution from the National Institute of Neurological Disorders and Stroke (NINDS). Additional support was provided through R01AG023629 from the National Institute on Aging (NIA). A full list of principal CHS investigators and institutions can be found at CHS-NHLBI.org. The provision of genotyping data was supported in part by the National Center for Advancing Translational Sciences, CTSI grant UL1TR001881, and the National Institute of Diabetes and Digestive and Kidney Disease Diabetes Research Center (DRC) grant DK063491 to the Southern California Diabetes Endocrinology Research Center. The content is solely the responsibility of the authors and does not necessarily represent the official views of the National Institutes of Health.

The China Atherosclerosis Study

This work was supported by the National Key R&D Program of China (2018YFE0115300, 2017YFC0908401,2017YFC0211703 and 2016YFC0206503) and the National Science Foundation of China (grants 81773537, 91857118, and 91439202).

China Kadoorie Biobank

China Kadoorie Biobank particularly acknowledges the participants, project staff in Beijing and Oxford and at the 10 regional CKB centers, and the China National Centre for Disease Control and Prevention (CDC) and its regional offices for assisting with the fieldwork. The CKB baseline survey and first re-survey were supported by the Kadoorie Charitable Foundation, Hong Kong. Long-term follow-up and data collection were supported by the Wellcome Trust, UK (212946/Z/18/Z, 202922/Z/16/Z, 104085/Z/14/Z, 088158/Z/09/Z), the National Key Research and Development Program of China (2016YFC0900500, 2016YFC0900501, 2016YFC0900504, 2016YFC1303904) and the National Natural Science Foundation of China (91846303, 81941018, 91843302, 81390540). DNA extraction and genotyping was supported by GlaxoSmithKline and the UK Medical Research Council (MC-PC-13049, MC-PC-14135). The project is supported by core funding to the Clinical Trial Service Unit and Epidemiological Studies Unit at Oxford University from the UK Medical Research Council (MC_UU_00017/1, MC_UU_12026/2, MC_U137686851), Cancer Research UK (C16077/A29186, C500/A16896) and the British Heart Foundation (CH/1996001/9454). Dr Holmes is supported by a British Heart Foundation Intermediate Clinical Research Fellowship (FS/18/23/33512).

China Health and Nutrition Survey

This research uses data from China Health and Nutrition Survey (CHNS). We are grateful to research grant funding from the National Institute for Health (NIH), the Eunice Kennedy Shriver National Institute of Child Health and Human Development (NICHD) for R01 HD30880, National Institute on Aging (NIA) for R01 AG065357, National Institute of Diabetes and Digestive and Kidney Diseases (NIDDK) for R01DK104371 and R01HL108427, the NIH Fogarty grant D43 TW009077 since 1989, and the China-Japan Friendship Hospital, Ministry of Health for support for CHNS 2009, Chinese National Human Genome Center at Shanghai since 2009, and Beijing Municipal Center for Disease Prevention and Control since 2011. We thank the National Institute for Nutrition and Health, China Center for Disease Control and Prevention, Beijing Municipal Center for Disease Control and Prevention, and the Chinese National Human Genome Center at Shanghai. Additional support for data analysis was provided by US NIH R01DK072193.

Cleveland Family Study

The CFS was supported by the National Institutes of Health, the National Heart, Lung, Blood Institute grants HL113338, R35HL13588, HL46380. BEC is supported by the NHLBI grant K01HL135405.

Cebu Longitudinal Health and Nutrition Survey

The Cebu Longitudinal Health and Nutrition Survey (CLHNS) was supported by US National Institutes of Health grants DK078150, TW005596, HL085144; pilot funds from RR020649, ES010126, and DK056350; and the Office of Population Studies Foundation in Cebu. Additional support for data analysis was provided by US NIH R01DK072193. Cassandra N Spracklen was supported by American Heart Association Postdoctoral Fellowships 15POST24470131 and 17POST33650016.

CoLaus study

The CoLaus study was and is supported by research grants from GlaxoSmithKline, the Faculty of Biology and Medicine of Lausanne, and the Swiss National Science Foundation (grants 33CSCO-122661, 33CS30-139468, 33CS30-148401 and 33CS30_177535/1).

Cooperative Health Research in South Tyrol

Full acknowledgements for the CHRIS study are reported at http://translational-medicine.biomedcentral.com/articles/10.1186/s12967-015-0704-9#Declarations. The CHRIS study was funded by the Department of Innovation, Research, and University of the Autonomous Province of Bolzano-South Tyrol.

CROATIA-Via and CROATIA-Korcula

The CROATIA_Vis and CROATIA_Korcula studies were funded by grants from the Medical Research Council (UK), European Commission Framework 6 project EUROSPAN (Contract No. LSHG-CT-2006-018947) and Republic of Croatia Ministry of Science, Education and Sports research grants. (108-1080315-0302). We would like to acknowledge the staff of several institutions in Croatia that supported the field work, including but not limited to The University of Split and Zagreb Medical Schools, Institute for Anthropological Research in Zagreb and Croatian Institute for Public Health.CH and VV are supported by an MRC University Unit Programme Grant MC_UU_00007/10 (QTL in Health and Disease)

The Danish study of Functional Disorders

This study was supported by TrygFonden (7-11-0213), the Lundbeck Foundation (R155-2013-14070), Novo Nordisk Foundation (NNF15OC0015896).

Diabetes Genetics Initiative of Broad Institute of Harvard and MIT, Lund University, and Novartis Institutes of BioMedical Research

The Botnia (DGI) study have been supported by grants from Folkhälsan Research Foundation, Sigrid Juselius Foundation, Ministry of Education, Nordic Center of Excellence in Disease Genetics, Gyllenberg Foundation, Swedish Cultural Foundation in Finland, Finnish Diabetes Research Foundation, Foundation for Life and Health in Finland, Finnish Medical Society, Paavo Nurmi Foundation, Perklén Foundation, Ollqvist Foundation, Närpes Health Care Foundation, the Municipal Health Care Center and Hospital in Jakobstad, Health Care Centers in Vasa, Närpes and Korsholm. This work was also partially supported by NIH grant R01-DK075787 to JNH. The skillful assistance of the Botnia Study Group is gratefully acknowledged

DIACORE

Cohort recruiting and management was funded by the KfH Stiftung Präventivmedizin e.V. (Carsten A. Böger). Genome-wide genotyping was funded the Else Kröner-Fresenius-Stiftung (2012_A147), the KfH Stiftung Präventivmedizin and the University Hospital Regensburg. The Deutsche Forschungsgemeinschaft (DFG, German Research Foundation) supported this work – Project-ID 387509280 – SFB 1350 (Subproject C6 to I.M.H.) and Iris Heid and Carsten Böger received funding by DFG BO 3815/4-1.

Electronic Medical Records and Genomics

eMERGE Network (Phase III): This phase of the eMERGE Network was initiated and funded by the NHGRI through the following grants: U01HG8657 (Group Health Cooperative/University of Washington); U01HG8685 (Brigham and Women’s Hospital); U01HG8672 (Vanderbilt University Medical Center); U01HG8666 (Cincinnati Children’s Hospital Medical Center); U01HG6379 (Mayo Clinic); U01HG8679 (Geisinger Clinic); U01HG8680 (Columbia University Health Sciences); U01HG8684 (Children’s Hospital of Philadelphia); U01HG8673 (Northwestern University); U01HG8701 (Vanderbilt University Medical Center serving as the Coordinating Center); U01HG8676 (Partners Healthcare/Broad Institute); and U01HG8664 (Baylor College of Medicine).

European Investigation into Cancer and Nutrition, Potsdam study

The recruitment phase of the EPIC-Potsdam Study was supported by the Federal Ministry of Science, Germany (01 EA 9401) and the European Union (SOC 95201408 05F02). The follow-up of the EPIC-Potsdam Study was supported by German Cancer Aid (70-2488-Ha I) and the European Community (SOC 98200769 05F02). This work was supported by a grant from the German Ministry of Education and Research (BMBF) and the State of Brandenburg (DZD grant 82DZD00302).

Family Heart Study

This work has been funded by the NIDDK grant R01-DK-089256, and the NHLBI grant R01HL117078

Fenland Study

The Fenland Study (10.22025/2017.10.101.00001) is funded by the Medical Research Council (MC_UU_12015/1). We are grateful to all the volunteers and to the General Practitioners and practice staff for assistance with recruitment. We thank the Fenland Study Investigators, Fenland Study Co-ordination team and the Epidemiology Field, Data and Laboratory teams. We further acknowledge support for genomics from the Medical Research Council (MC_PC_13046).

Finland-United States Investigation of NIDDM Genetics

Support for FUSION was provided by NIH grants R01-DK062370, R01-DK072193, and intramural project number 1Z01-HG000024. Genome-wide genotyping was conducted by the Johns Hopkins University Genetic Resources Core Facility SNP Center at the Center for Inherited Disease Research (CIDR), with support from CIDR NIH contract no. N01-HG-65403.

Finnish Twin Cohort

Phenotype and genotype data collection in the twin cohort has been supported by the Wellcome Trust Sanger Institute, the Broad Institute, ENGAGE – European Network for Genetic and Genomic Epidemiology, FP7-HEALTH-F4-2007, grant agreement number 201413, National Institute of Alcohol Abuse and Alcoholism (grants AA-12502, AA-00145, and AA-09203 to Richard J Rose and AA15416 and K02AA018755 to Danielle M Dick) and the Academy of Finland (grants 100499, 205585, 118555, 141054, 264146, 308248, and 312073 to JKaprio). JKaprio acknowledges support by the Academy of Finland (grants 265240, 263278).

FINRISK

FINRISK has been mainly funded from budgetary funds of the Finnish Institute for Health and Welfare. Additional funds have been obtained from domestic foundations. VS has been supported by the Finnish Foundation for Cardiovascular Research.

Framingham Heart Study

This research was conducted in part using data and resources from the Framingham Heart Study of the National Heart Lung and Blood Institute of the National Institutes of Health and Boston University School of Medicine. The analyses reflect intellectual input and resource development from the Framingham Heart Study investigators participating in the SNP Health Association Resource (SHARe) project. This work was partially supported by the National Heart, Lung and Blood Institute's Framingham Heart Study (Contract Nos. N01-HC-25195, HHSN268201500001I and 75N92019D00031) and its contract with Affymetrix, Inc for genotyping services (Contract No. N02-HL-6-4278).

Genetic Park of Cilento and Vallo di Diano Project

This work was supported by IDF SHARID grant from the Italian Ministry of Universities and Research

GENIE_UK

Original genotype data was generated as part of the GEnetics of Nephropathy an International Effort (GENIE) consortium through the Medical Research Council (MC_PC_15025) and the Public Health Agency R&D Division (STL/4760/13), Science Foundation Ireland (SFI15/US/B3130) and NIH R01_DK081923 and R01_DK105154. Science Foundation Ireland and the Department for the Economy, Northern Ireland US partnership award 15/IA/3152. The GENIE_UK data includes samples recruited as part of the Warren3/U.K. GoKinD Study Group, jointly funded by Diabetes UK and the Juvenile Diabetes Research Foundation and includes the following individuals: Professor A.P. Maxwell, Prof A.J. McKnight, Dr. D.A. Savage (Belfast); Dr. J. Walker (Edinburgh); Dr. S. Thomas, Professor G.C. Viberti (London); Professor A.J.M. Boulton (Manchester); Professor S. Marshall (Newcastle); Professor A.G. Demaine and Dr. B.A. Millward (Plymouth); and Professor S.C. Bain (Swansea).

German Chronic Kidney Disease Study

The GCKD study is supported by the German Ministry of Education and Research (Bundesministerium für Bildung und Forschung, FKZ 01ER 0804, 01ER 0818, 01ER 0819, 01ER 0820, and 01ER 0821) and the KfH Foundation for Preventive Medicine (Kuratorium für Heimdialyse und Nierentransplantation e.V.–Stiftung Präventiv-medizin) and corporate sponsors (www.gckd.org). Furthermore, this study was partially funded by the H2020-IMI2 consortium BEAt-DKD (Biomarker Enterprise to Attack Diabetic Kidney Disease; grant number: 115974). Whole-genome SNP microarray genotyping in the GCKD study was supported by Bayer Pharma Aktiengesellschaft (AG).

Genetic Epidemiology Network of Arteriopathy

Support for the Genetic Epidemiology Network of Arteriopathy (GENOA) was provided by the

National Heart, Lung and Blood Institute (HL054464, HL054457, HL054481, HL100185,

and HL119443) of the National Institutes of Health. We would also like to thank the families that

participated in the GENOA study.

German MI Family Study

Funding was received from the German Federal Ministry of Education and Research (BMBF) in the context of the e:Med program (e:AtheroSysMed and sysINFLAME), the FP7 European Union project CVgenes@target (261123) and a grant from the Fondation Leducq (CADgenomics: Understanding Coronary Artery Disease Genes, 12CVD02).

Gene-Lifestyle Interactions and Complex Traits Involved in Elevated Disease Risk-V2

European Research Council: CoG-2015_681742_NASCENT; Swedish Research Council; Novo Nordisk Foundation; European Diabetes Research Foundation; Swedish Heart Lung Foundation

Generation Scotland

Generation Scotland received core support from the Chief Scientist Office of the Scottish Government Health Directorates [CZD/16/6] and the Scottish Funding Council [HR03006] and is currently supported by the Wellcome Trust [216767/Z/19/Z]. Genotyping of the GS:SFHS samples was carried out by the Genetics Core Laboratory at the Edinburgh Clinical Research Facility, University of Edinburgh, Scotland and was funded by the Medical Research Council UK and the Wellcome Trust (Wellcome Trust Strategic Award “STratifying Resilience and Depression Longitudinally” (STRADL) Reference 104036/Z/14/Z).

Genetic Overlap between Metabolic and Psychiatric traits/TEENAGE

This research has been co-financed by the European Union (European Social Fund—ESF) and Greek national funds through the Operational Program “Education and Lifelong Learning” of the National Strategic Reference Framework (NSRF)—Research Funding Program: Heracleitus II. Investing in knowledge society through the European Social Fund. This work was funded by the Wellcome Trust (098051). We thank all study participants and their families as well as all volunteers for their contribution in this study. We thank the Sample Management and Genotyping Facilities staff at the Wellcome Trust Sanger Institute for sample preparation, quality control and genotyping. We are grateful to Georgia Markou, Laiko General Hospital Diabetes Centre, Maria Emetsidou and Panagiota Fotinopoulou, Hippokratio General Hospital Diabetes Centre, Athina Karabela, Dafni Psychiatric Hospital, Eirini Glezou and Marios Mangioros, Dromokaiteio Psychiatric Hospital, Angela Rentari, Harokopio University of Athens, and Danielle Walker, Wellcome Trust Sanger Institute.

Genetic Regulation of Arterial Pressure In humans in the Community

NJS holds an NIHR Senior Investigator award.

Genomic Research Cohorts for CCMB Diabetes Study

The work was supported by funding from Council of Scientific Industrial Research (CSIR), Ministry of Science and Technology, Government of India, New Delhi, INDIA and forms part of Network Project funds. We acknowledge the contribution of all the participants under the study and also of the members of GRC-Group at CSIR-Centre for Cellular and Molecular Biology, Hyderabad. INDIA. MA was supported by funds from Council of Scientific Industrial Research (CSIR), Ministry of Science and Technology, Government of India, New Delhi, INDIA.

GoDARTS

We are grateful to all the participants in this study, the general practitioners, the Scottish School of Primary Care for their help in recruiting the participants, and to the whole team, which includes interviewers, computer and laboratory technicians, clerical workers, research scientists, volunteers, managers, receptionists, and nurses. The study complies with the Declaration of Helsinki. We acknowledge the support of the Health Informatics Centre, University of Dundee for managing and supplying the anonymised data and NHS Tayside, the original data owner. The Wellcome Trust United Kingdom Type 2 Diabetes Case Control Collection (GoDARTS) was funded by The Wellcome Trust (072960/Z/03/Z, 084726/Z/08/Z, 084727/Z/08/Z, 085475/Z/08/Z, 085475/B/08/Z). MMcC was a Wellcome Investigator and NIHR Senior Investigator. This work was supported by: NIDDK (U01-DK105535) and Wellcome (090532, 098381, 106130, 203141, 212259)

Genetics of Lipid Lowering and Diet Network (GOLDN)

GOLDN biospecimens, baseline phenotype data, and intervention phenotype data were collected with funding from National Heart, Lung and Blood Institute (NHLBI) grant U01 HL072524. Whole-genome sequencing in GOLDN was funded by NHLBI grant R01 HL104135-04S1.

Greek Recurrent Myocardial Infarction Cohort /NAFLD Study

Work was supported by the British Heart Foundation (BHF) grant RG/14/5/30893 (P.D.) and forms part of the research themes contributing to the translational research portfolios of the Barts Biomedical Research Centre funded by the UK National Institute for Health Research (NIHR). O.G. has received funding from the British Heart Foundation (BHF) (FS/14/66/3129)

Healthy Aging in Neighborhoods of Diversity across the Life Span

National Institute on Aging NIH AG-000513

Health 2006,2008,2010

The Health2006 was financially supported by grants from the Velux Foundation; The Danish Medical Research Council, Danish Agency for Science, Technology and Innovation; The Aase and Ejner Danielsens Foundation; ALK-Abello A/S, Hørsholm, Denmark, and Research Centre for Prevention and Health, the Capital Region of Denmark. Health2008 was supported by the Timber Merchant Vilhelm Bang’s Foundation, the Danish Heart Foundation (Grant number 07-10-R61-A1754-B838-22392F), and the Health Insurance Foundation (Helsefonden) (Grant number 2012B233).

Hellenic Isolated Cohorts MANOLIS and POMAK

This work was funded by the Wellcome Trust (098051) and the European Research Council (ERC-2011-StG 280559-SEPI). The MANOLIS cohort is named in honour of Manolis Giannakakis, 1978-2010. We thank the residents of the Mylopotamos villages for taking part.

Hein Nixdorf Recall Study

This work was supported by the Heinz Nixdorf Foundation; the German Research Council [projects SI 236/8-1, SI 236/9-1, ER 155/6-1, ER 155/6-2].

Health and Retirement Study

HRS is supported by the National Institute on Aging (NIA U01AG009740). The genotyping was

funded separately by the National Institute on Aging (RC2 AG036495, RC4 AG039029). Our

genotyping was conducted by the NIH Center for Inherited Disease Research (CIDR) at Johns

Hopkins University. Genotyping quality control and final preparation of the data were

performed by the Genetics Coordinating Center at the University of Washington and the School

of Public Health at the University of Michigan.

Hispanic Community Health Study/Study of Latinos

The baseline examination of HCHS/SOL was carried out as a collaborative study supported by contracts from the NHLBI to the University of North Carolina (N01-HC65233), University of Miami (N01-HC65234), Albert Einstein College of Medicine (N01-HC65235), Northwestern University (N01-HC65236), and San Diego State University (N01-HC65237). The following institutes, centers, and offices contributed to the first phase of HCHS/SOL through a transfer of funds to the NHLBI: National Institute on Minority Health and Health Disparities, National Institute on Deafness and Other Communication Disorders, National Institute of Dental and Craniofacial Research (NIDCR), National Institute of Diabetes and Digestive and Kidney Diseases (NIDDK), National Institute of Neurological Disorders and Stroke, and NIH Office of Dietary Supplements. The Genetic Analysis Center at the University of Washington was supported by NHLBI and NIDCR contracts (HHSN268201300005C AM03 and MOD03). Additional analysis support was provided by NIDDK grant 1R01DK101855-01, NHLBI grant N01HC65233, and AHA grant 13GRNT16490017. Genotyping efforts were supported by the NIH Department of Health and Human Services (HSN26220/20054C), National Center for Advancing Translational Science Clinical Translational Science Institute (UL1TR000124), and NIDDK Diabetes Research Center (DK063491). Kristin L. Young was supported by R21 HL140419. Heather M Highland was funded by NHLBI training grant T32 HL007055, T32 HL129982, ADA Grant #1-19-PDF-045, and R01HL142825.

Hunter Community Study

The authors would like to thank the men and women participating in the HCS as well as all the staff, investigators and collaborators who have supported or been involved in the project to date. The cohort was supported by grants from the University of Newcastle Strategic Initiatives Fund, the Gladys M Brawn Senior Research Fellowship scheme; and the Vincent Fairfax Family Foundation, a private philanthropic trust. The Hunter Medical Research Institute provided media support during the initial recruitment of participants; and Dr Anne Crotty and Prof. Levi provided financial support towards freezing costs for the long-term storage of participant blood samples.

HUNT Study

The HUNT Study is a collaboration between the HUNT Research Centre (Faculty of Medicine and Health Sciences, NTNU, Norwegian University of Science and Technology), Nord-Trøndelag County Council, Central Norway Regional Health Authority, and the Norwegian Institute of Public Health. The genotyping in HUNT was financed by the National Institutes of Health; University of Michigan; the Research Council of Norway; the Liaison Committee for Education, Research and Innovation in Central Norway; and the Joint Research Committee between St Olavs hospital and the Faculty of Medicine and Health Sciences, NTNU.

Hypertension Genetic Epidemiology Network (HyperGEN + HyperGEN LVH)

The Hypertension Genetic Epidemiology Network (HyperGEN) Study is part of the National Heart, Lung, and Blood Institute (NHLBI) Family Blood Pressure Program; collection of the data represented here was supported by grants U01 HL054472, U01 HL054473, U01 HL054495, and U01 HL054509. The HyperGEN: Genetics of Left Ventricular Hypertrophy Study was supported by NHLBI grant R01 HL055673 with whole-genome sequencing made possible by supplement -18S1.

Hypertension Genetic Epidemiology Network (Axiom chip)

The study was support by the National Institutes of Health, the National Heart, Lung, Blood Institute grant HL086718

INdian DIabetes COnsortium

The study was majorly supported by Council of Scientific and Industrial Research (CSIR), Government of India through CARDIOMED project Grant Number: BSC0122 provided to CSIR-Institute of Genomics and Integrative Biology. Study was also partially funded by Department of Science and Technology-PURSE-II (DST/SR/PURSE II/11) given to Jawaharlal Nehru University. We are very much thankful to the INdian DIabetes COnsortium (INDICO) and all the study participants who have participated in the study.

Inter99

The Inter99 was initiated by Torben Jørgensen (PI), Knut Borch-Johnsen (co-PI), Hans Ibsen and Troels F. Thomsen. The steering committee comprises the former two and Charlotta Pisinger. The study was financially supported by research grants from the Danish Research Council, the Danish Centre for Health Technology Assessment, Novo Nordisk Inc., Research Foundation of Copenhagen County, Ministry of Internal Affairs and Health, the Danish Heart Foundation, the Danish Pharmaceutical Association, the Augustinus Foundation, the Ib Henriksen Foundation, the Becket Foundation, and the Danish Diabetes Association. Novo Nordisk Foundation Center for Basic Metabolic Research is an independent Research Center, based at the University of Copenhagen, Denmark and partially funded by an unconditional donation from the Novo Nordisk Foundation (www.cbmr.ku.dk) (Grant number NNF18CC0034900).

INGI-FVG: This research was funded by D70-RESRICGIROTTO to Giorgia Girotto, and Italian Ministry of Health—RC 01/21 to Maria Pina Concas. We thank all the inhabitants of FVG who participated in the projects

Jackson Heart Study

The Jackson Heart Study (JHS) is supported and conducted in collaboration with Jackson State University (HHSN268201800013I), Tougaloo College (HHSN268201800014I), the Mississippi State Department of Health (HHSN268201800015I) and the University of Mississippi Medical Center (HHSN268201800010I, HHSN268201800011I and HHSN268201800012I) contracts from the National Heart, Lung, and Blood Institute (NHLBI) and the National Institute on Minority Health and Health Disparities (NIMHD). The authors also wish to thank the staffs and participants of the JHS.

The views expressed in this manuscript are those of the authors and do not necessarily represent the views of the National Heart, Lung, and Blood Institute; the National Institutes of Health; or the U.S. Department of Health and Human Services.

The Johnston County Osteoarthritis Project

The JoCoOA is supported in part by S043, S1734, & S3486 from the CDC/Association of Schools of Public Health; 5-P60-AR30701 & 5-P60-AR49465 from NIAMS/NIH, and U01 DP003206 from the CDC

Japan PGx Data Science Consortium

The authors thank the Japan PGx Data Science Consortium (JPDSC) for kindly providing genotype and phenotype data. The JPDSC was comprised of six pharmaceutical companies in Japan, namely Astellas Pharma, Inc.; Daiichi Sankyo Co., Ltd.; Mitsubishi Tanabe Pharma Corporation; Otsuka Pharmaceutical Co., Ltd.; Taisho Pharmaceutical Co., Ltd.; and Takeda Pharmaceutical Co., Ltd.

Korea Association REsource

The KARE cohort was supported by grants from Korea Centers for Disease Control and Prevention (4845–301, 4851–302, 4851–307) and an intramural grant from the Korea National Institute of Health (2019-NG-053-01).

KORA

The KORA study was initiated and financed by the Helmholtz Zentrum München – German Research Center for Environmental Health, which is funded by the German Federal Ministry of Education and Research (BMBF) and by the State of Bavaria. KORA research has been supported within the Munich Center of Health Sciences (MC-Health), Ludwig-Maximilians-Universität, as part of LMUinnovativ and is funded by the Bavarian State Ministry of Health and Care through the research project DigiMed Bayern (www.digimed-bayern.de). The German Center for Diabetes Research (DZD) was supported by the German Federal Ministry of Education and Research (BMBF).

Korea National Diabetes Program

This study was supported by a grant from the Korea Healthcare Technology R&D Project, Ministry of Health and Welfare Republic of Korea (HI10C2020).

Kangbuk Samsung Cohort Study (KSCS)

This KSCS research was supported by the National Research Foundation of Korea (NRF) grant funded by the Korea government (MSIT) (NRF-2020R1A2C1012931) and by the Medical Research Funds from Kangbuk Samsung Hospital. We are thankful for the computing resources provided by the Global Science experimental Data hub Center (GSDC) Project and the Korea Research Environment Open NETwork (KREONET) of the Korea Institute of Science and Technology Information (KISTI).

Kuwait Obesity Diabetes Genetic Program

Kuwait Foundation for Advancement of Sciences is acknowledged for institutional funding.

Leicester BIORESOURCE

CPN is funded by the British Heart Foundation. NJS holds an NIHR Senior Investigator award.

Long Life Family Study

This work has been funded by the NIA grants, U01AG023746, U01AG023712, U01AG023749, U01AG023755, U01AG023744, and U19AG063893

London Life Sciences Prespective Population Study

The LOLIPOP study is supported by the National Institute for Health Research (NIHR) Comprehensive Biomedical Research Centre Imperial College Healthcare NHS Trust, the British Heart Foundation (SP/04/002), the Medical Research Council (G0601966, G0700931), the Wellcome Trust (084723/Z/08/Z, 090532 & 098381) the NIHR (RP-PG-0407-10371), the NIHR Official Development Assistance (ODA, award 16/136/68), the European Union FP7 (EpiMigrant, 279143) and H2020 programs (iHealth-T2D, 643774). We acknowledge support of the MRC-PHE Centre for Environment and Health, and the NIHR Health Protection Research Unit on Health Impact of Environmental Hazards. The work was carried out in part at the NIHR/Wellcome Trust Imperial Clinical Research Facility. The views expressed are those of the author(s) and not necessarily those of the Imperial College Healthcare NHS Trust, the NHS, the NIHR or the Department of Health. We thank the participants and research staff who made the study possible. JC is supported by the Singapore Ministry of Health’s National Medical Research Council under its Singapore Translational Research Investigator (STaR) Award (NMRC/STaR/0028/2017).

Ludwigshafen Risk and Cardiovascular Health Study

The LURIC study was supported by the 7th Framework Program (AtheroRemo, grant agreement number 201668 and RiskyCAD, grant agreement number 305739) of the European Union, the INTERREG IV Oberrhein Program (Project A28) with support from the European Regional Development Fund (ERDF) and the Wissenschaftsoffensive TMO, and European Union’s Horizon 2020 research and innovation programme under the ERA-Net Cofund action N° 727565 (OCTOPUS project) and the German Ministry of Education and Research (grant number 01EA1801A)

Metabolic Syndrome in Men

The METSIM study was funded by the Academy of Finland (grants no. 77299 and 124243).

Mexico City Study

This study was supported in Mexico by the Fondo Sectorial de Investigación en Salud y Seguridad Social (SSA/IMSS/ISSSTECONACYT, project 150352), Temas Prioritarios de Salud Instituto Mexicano del Seguro Social (2014-FIS/IMSS/PROT/PRIO/14/34). The authors would like to thank Araceli Méndez Padrón for technical support. In Canada, this research was enabled in part by support provided by Compute Ontario (www.computeontario.ca), WestGrid (www.westgrid.ca) and Compute Canada (www.compute.canada.ca).

Million Veteran Program (MVP)

This research is supported by funding from the Department of Veterans Affairs Office of Research and Development, Million Veteran Program Grant I01-BX003340 and I01-BX003362. This publication does not represent the views of the Department of Veterans Affairs or the United States Government. A list of MVP investigators can be found in supplementary materials.

Montreal Heart Institute Biobank

We thank all participants and staff of the André and France Desmarais MHI Biobank. This work was funded by the Canadian Institutes of Health Research (PJT #156248), the Canada Research Chair Program, Genome Quebec and Genome Canada, and the Montreal Heart Institute Foundation.

Mount Sinai BioMe Biobank

The Mount Sinai BioMe Biobank has been supported by The Andrea and Charles Bronfman Philanthropies and in part by Federal funds from the NHLBI and NHGRI (U01HG00638001; U01HG007417; X01HL134588). We thank all participants in the Mount Sinai Biobank. We also thank all our recruiters who have assisted and continue to assist in data collection and management and are grateful for the computational resources and staff expertise provided by Scientific Computing at the Icahn School of Medicine at Mount Sinai.

MRC National Survey of Health and Development

We thank NSHD study members who took part in the data collections for their continuing support. This work was supported by the UK Medical Research Council which provides core funding for the MRC National Survey of Health and Development (MC_UU_12019/06).

Multi-Ethnic Study of Atherosclerosis

MESA and the MESA SHARe projects are conducted and supported by the National Heart, Lung, and Blood Institute (NHLBI) in collaboration with MESA investigators. Support for MESA is provided by contracts 75N92020D00001, HHSN268201500003I, N01-HC-95159, 75N92020D00005, N01-HC-95160, 75N92020D00002, N01-HC-95161, 75N92020D00003, N01-HC-95162, 75N92020D00006, N01-HC-95163, 75N92020D00004, N01-HC-95164, 75N92020D00007, N01-HC-95165, N01-HC-95166, N01-HC-95167, N01-HC-95168, N01-HC-95169, UL1-TR-000040, UL1-TR-001079, UL1-TR-001420. Also supported in part by the National Center for Advancing Translational Sciences, CTSI grant UL1TR001881, and the National Institute of Diabetes and Digestive and Kidney Disease Diabetes Research Center (DRC) grant DK063491 to the Southern California Diabetes Endocrinology Research Center.

Nutrition and Health of Aging Population in China

This work was supported by the Major Project of the Ministry of Science and Technology of China (2017YFC0909701, 2016YFC1304903), the National Natural Science Foundation of China (81970684, 81700700), the Chinese Academy of Sciences (ZDBS-SSW-DQC-02, KSCX2-EW-R-10, KJZD-EW-L14-2-2) and Shanghai Municipal Science and Technology Major Project (2017SHZDZX01)

the Nagahama Prospective Cohort for Comprehensive Human Bioscience (the Nagahama Study)

This work was supported by a university grant, Center of Innovation Program, Global University Project, and a Grant-in-Aid for Scientific Research (25293141, 26670313, 26293198, 17H04182, 17H04126, 17H04123, 18K18450) from the Ministry of Education, Culture, Sports, Science and Technology of Japan; Practical Research Project for Rare/Intractable Diseases (ek0109070, ek0109283, ek0109196, ek0109348), Research and Development Grants for Dementia (dk0207006, dk0207027), Program for an Integrated Database of Clinical and Genomic Information (kk0205008), Practical Research Project for Lifestyle-related Diseases including Cardiovascular Diseases and Diabetes Mellitus (ek0210066, ek0210096, ek0210116), Research Program for Health Behavior Modification by Utilizing IoT (le0110005, le0110013), and Research and Development Grants for Longevity Science (dk0110040) from the Japan Agency for Medical Research and Development (AMED); Takeda Medical Research Foundation, Mitsubishi Foundation, Daiwa Securities Health Foundation, and Sumitomo Foundation.

Nijmegen Biomedical Study

The Nijmegen Biomedical Study is a population-based survey conducted at the Department for Health Evidence and the Department of Laboratory Medicine of the Radboud university medical center. Principal investigators of the Nijmegen Biomedical Study are L.A.L.M. Kiemeney, A.L.M. Verbeek, D.W. Swinkels en B. Franke

The Netherlands Epidemiology of Obesity Study

The authors of the NEO study thank all individuals who participated in the Netherlands Epidemiology in Obesity study, all participating general practitioners for inviting eligible participants and all research nurses for collection of the data. We thank the NEO study group, Pat van Beelen, Petra Noordijk and Ingeborg de Jonge for the coordination, lab and data management of the NEO study. The genotyping in the NEO study was supported by the Centre National de Génotypage (Paris, France), headed by Jean-Francois Deleuze. The NEO study is supported by the participating Departments, the Division and the Board of Directors of the Leiden University Medical Center, and by the Leiden University, Research Profile Area Vascular and Regenerative Medicine. Dennis Mook-Kanamori is supported by Dutch Science Organization (ZonMW-VENI Grant 916.14.023).

Northern Ireland Cohort for Longitudinal Study of Ageing

The analysis of molecular biomarkers for NICOLA’s Wave 1 was funded by the Economic and Social Research Council, award reference ES/L008459/1. Generic analysis of data was supported by ESRC (ES/L008459/1) and the Science Foundation Ireland-Department for the Economy (SFI-DfE) Investigator Program Partnership Award (15/IA/3152). We are grateful to all the participants of the NICOLA Study, and the whole NICOLA team, which includes nursing staff, research scientists, clerical staff, computer and laboratory technicians, managers and receptionists. The Atlantic Philanthropies, the Economic and Social Research Council, the UKCRC Centre of Excellence for Public Health Northern Ireland, the Centre for Ageing Research and Development in Ireland, the Office of the First Minister and Deputy First Minister, the Health and Social Care Research and Development Division of the Public Health Agency, the Wellcome Trust/Wolfson Foundation and Queen’s University Belfast provide core financial support for NICOLA. The authors alone are responsible for the interpretation of the data and any views or opinions presented are solely those of the authors and do not necessarily represent those of the NICOLA Study team.

Netherlands Twin Register

Funding was obtained from the Netherlands Organization for Scientific Research (NWO) and The Netherlands Organisation for Health Research and Development (ZonMW) grants 904-61-090, 985-10-002, 904-61-193,480-04-004, 400-05-717, Addiction-31160008, 016-115-035, 400-07-080, Middelgroot-911-09-032, NWO-Groot 480-15-001/674, Center for Medical Systems Biology (CSMB, NWO Genomics), NBIC/BioAssist/RK(2008.024), Biobanking and Biomolecular Resources Research Infrastructure (BBMRI –NL, 184.021.007 and 184.033.111), X-Omics 184-034-019; Spinozapremie (NWO- 56-464-14192), KNAW Academy Professor Award (PAH/6635) and University Research Fellow grant (URF) to DIB; Amsterdam Public Health research institute (former EMGO+) , Neuroscience Amsterdam research institute (former NCA) ; the European Community's Fifth and Seventh Framework Program (FP5- LIFE QUALITY-CT-2002-2006, FP7- HEALTH-F4-2007-2013, grant 01254: GenomEUtwin, grant 01413: ENGAGE ; the European Research Council (ERC Starting 284167, ERC Consolidator 771057, ERC Advanced 230374), Rutgers University Cell and DNA Repository (NIMH U24 MH068457-06), the National Institutes of Health (NIH, R01D0042157-01A1, MH081802, DA018673, R01 DK092127-04, Grand Opportunity grants 1RC2 MH089951); the Avera Institute for Human Genetics, Sioux Falls, South Dakota (USA). Part of the genotyping and analyses were funded by the Genetic Association Information Network (GAIN) of the Foundation for the National Institutes of Health. Computing was supported by NWO through grant 2018/EW/00408559, BiG Grid, the Dutch e-Science Grid and SURFSARA.

Ogliastra Genetic Park

The study was supported by grant from the Italian Ministry of Education, University and Research (MIUR) n°: 5571/DSPAR/2002. We express our gratitude to all the study participants for their contributions and to the municipal administrations for their economic and logistic support.

Orkney Complex Disease Study

The Orkney Complex Disease Study (ORCADES) was supported by the Chief Scientist Office of the Scottish Government (CZB/4/276, CZB/4/710), a Royal Society URF to J.F.W., the MRC Human Genetics Unit quinquennial programme “QTL in Health and Disease”, Arthritis Research UK and the European Union framework program 6 EUROSPAN project (contract no. LSHG-CT-2006-018947). DNA extractions were performed at the Edinburgh Clinical Research Facility, University of Edinburgh. We would like to acknowledge the invaluable contributions of the research nurses in Orkney, the administrative team in Edinburgh and the people of Orkney.

Osteoporosis Fractures in Men, The Gothenburg Study

MrOS in Sweden is supported by the Swedish Research Council, the Swedish Foundation for Strategic Research, the ALF/LUA research grant in Gothenburg, the Lundberg Foundation, the Knut and Alice Wallenberg Foundation, the Torsten Soderberg Foundation, and the Novo Nordisk Foundation.

Oxford Biobank

This work was supported by the British Heart Foundation (RG/17/1/32663)

Penn Medicine BioBank

We acknowledge the Penn Medicine BioBank (PMBB) for providing data and thank the patient-participants of Penn Medicine who consented to participate in this research program. We would also like to thank the Penn Medicine BioBank team and Regeneron Genetics Center for providing genetic variant data for analysis. The PMBB is approved under IRB protocol# 813913 and supported by Perelman School of Medicine at University of Pennsylvania

pcosN

YSC is supported by by the National Research Foundation of Korea (NRF) Grant funded by the Ministry of Education (NRF-2020R1I1A2075302) and Hallym University Research Fund 2020 (HRF-202002-008).

PRecOcious Coronary ARtery DISease

PROCARDIS was supported by the European Community Sixth Framework Program (LSHM-CT- 2007-037273), AstraZeneca, the Swedish Research Council, the Knut and Alice Wallenberg Foundation, the Swedish Heart-Lung Foundation, the Torsten and Ragnar Soderberg Foundation, the Strategic Cardiovascular Program of Karolinska Institutet and Stockholm County Council, the Foundation for Strategic Research and the Stockholm County Council (560283). HW is supported by the British Heart Foundation Centre for Research Excellence, Wellcome Trust core award (090532/Z/09/Z, 203141/Z/16/Z, 201543/B/16/Z), European Union Seventh Framework Programme FP7/2007-2013 under grant agreement no. HEALTH-F2-2013-601456 (CVGenes@Target) & the TriPartite Immunometabolism Consortium [TrIC]-Novo Nordisk Foundation’s Grant number NNF15CC0018486. AG acknowledge support from the Wellcome Trust (201543/B/16/Z), European Union Seventh Framework Programme FP7/2007-2013 under grant agreement no. HEALTH-F2-2013-601456 (CVGenes@Target) & the TriPartite Immunometabolism Consortium [TrIC]-Novo Nordisk Foundation’s Grant number NNF15CC0018486. Maria Sabater-Lleal is supported by a Miguel Servet contract from the ISCIII Spanish Health Institute (CP17/00142) and co-financed by the European Social Fund.

Quebec Family Study

The Quebec Family Study was funded by multiple grants from the Medical Research Council of Canada and the Canadian Institutes of Health Research. This work was supported by a team grant from the Canadian Institutes of Health Research (FRN-CCT-83028)

QIMR Berghofer Genetic Epidemiology Cohort

Funded by the Australian National Health and Medical Research Council (241944, 339462, 389927, 389875, 389891, 389892, 389938, 442915, 442981, 496739, 552485, 552498), the Australian Research Council (A7960034, A79906588, A79801419, DP0770096, DP0212016, DP0343921) and the U.S. National Institutes of Health (AA07535, AA10248, AA11998, AA13320, AA13321, AA13326, AA14041, AA17688, DA12854, MH66206).

The Raine Study

The Raine Study was supported by the National Health and Medical Research Council of Australia [grant numbers 572613, 403981, and 1059711] and the Canadian Institutes of Health Research [grant number MOP-82893]. The authors are grateful to the Raine Study participants and their families, and to the Raine Study team for cohort coordination and data collection. The authors gratefully acknowledge the NHMRC for their long term funding to the study over the last 30 years and also the following institutes for providing funding for Core Management of the Raine Study: The University of Western Australia (UWA), Curtin University, Women and Infants Research Foundation, Telethon Kids Institute, Edith Cowan University, Murdoch University, The University of Notre Dame Australia and The Raine Medical Research Foundation. This work was supported by resources provided by the Pawsey Supercomputing Centre with funding from the Australian Government and Government of Western Australia.

Relationship between Insulin Sensitivity and Cardiovascular Disease

The RISC Study is partly supported by EU grant QLG1-CT-2001-01252.

Religious Orders Study & the Rush Memory and Aging Project

P30AG10161, R01AG15819, R10AG17917, U01AG46152, U01AG61356 and the Translational Genomics Research Institute

Rotterdam Study 1,2,3

The generation and management of GWAS genotype data for the Rotterdam Study (RS I, RS II, RS III) was executed by the Human Genotyping Facility of the Genetic Laboratory of the Department of Internal Medicine, Erasmus MC, Rotterdam, The Netherlands. The GWAS datasets are supported by the Netherlands Organisation of Scientific Research NWO Investments (nr. 175.010.2005.011, 911-03-012), the Genetic Laboratory of the Department of Internal Medicine, Erasmus MC, the Research Institute for Diseases in the Elderly (014-93-015; RIDE2), the Netherlands Genomics Initiative (NGI)/Netherlands Organisation for Scientific Research (NWO) Netherlands Consortium for Healthy Aging (NCHA), project nr. 050-060-810. We thank Pascal Arp, Mila Jhamai, Marijn Verkerk, Lizbeth Herrera and Marjolein Peters, PhD, and Carolina Medina-Gomez, PhD, for their help in creating the GWAS database, and Linda Broer,PhD, and Carolina Medina-Gomez, PhD, for the creation and analysis of imputed data. The Rotterdam Study is funded by Erasmus Medical Center and Erasmus University, Rotterdam, Netherlands Organization for the Health Research and Development (ZonMw), the Research Institute for Diseases in the Elderly (RIDE), the Ministry of Education, Culture and Science, the Ministry for Health, Welfare and Sports, the European Commission (DG XII), and the Municipality of Rotterdam. The authors are very grateful to the study participants, the staff from the Rotterdam Study (particularly L. Buist and J.H. van den Boogert) and the participating general practitioners and pharmacists.

Sardinia

Supported by contracts (HHSN271201600005C, to F. Cucca) from the Intramural Research Program of the 366 National Institute on Aging, National Institutes of Health (NIH)

Shanghai Breast Cancer Study

The generation and management of GWAS genotype data for the SBCS was supported by R01CA64277 and R01CA15847.

Steno Diabetes Center T2D Cases

Novo Nordisk Foundation Center for Basic Metabolic Research is an independent Research Center, based at the University of Copenhagen, Denmark and partially funded by an unconditional donation from the Novo Nordisk Foundation (www.cbmr.ku.dk) (Grant number NNF18CC0034900).

Study of Health in Pomerania

SHIP (Study of Health in Pomerania) and SHIP-TREND both represent population-based studies. SHIP is supported by the German Federal Ministry of Education and Research (Bundesministerium für Bildung und Forschung (BMBF); grants 01ZZ9603, 01ZZ0103, and 01ZZ0403) and the German Research Foundation (Deutsche Forschungsgemeinschaft (DFG); grant GR 1912/5-1). SHIP and SHIP-TREND are part of the Community Medicine Research net (CMR) of the University of Greifswald which is funded by the BMBF as well as the Ministry for Education, Science and Culture and the Ministry of Labor, Equal Opportunities, and Social Affairs of the Federal State of Mecklenburg-West Pomerania. The CMR encompasses several research projects that share data from SHIP.

SIGMA1,2

This work was conducted as part of the Slim Initiative in Genomic Medicine for the Americas (SIGMA), a joint U.S.-Mexico project funded by the Carlos Slim Foundation. The Universidad Nacional Autónoma de México/Instituto Nacional de Ciencias Médicas and Nutrición Salvador Zubirán (UNAM/INCMNSZ) diabetes study was supported by the Consejo Nacional de Ciencia y Tecnología (grants 138826, 128877, and CONACyT-SALUD 2009-01-115250) and a grant from Dirección General de Asuntos del Personal Académico (UNAM, IT 214711). The Diabetes in Mexico Study (DMS) was supported by the Consejo Nacional de Ciencia y Tecnología (grant 86867) and by the Carlos Slim Foundation. The Mexico City Diabetes Study (MCDS) was supported by National Institutes of Health (grant R01HL24799 from the National Heart, Lung, and Blood Institute) and by the Consejo Nacional de Ciencia y Tecnología (grants 2092, M9303, F677-M9407, 251M, 2005-C01-14502, and SALUD 2010-2- 151165). The Multiethnic Cohort Study of Diet and Cancer (MEC) was supported by National Institutes of Health (grants CA54281 and CA063464). This project was also supported by an American Diabetes Association Innovative and Clinical Translational Award 1-19-ICTS-068 and by DGAPA/UNAM Funding Project IN215219.

Singapore Eye Epidemiology Diseases (SEED)

Singapore Chinese Eye Study (SCES) + Singapore Malay Eye Study (SiMES) + Singapore Indian Eye Study (SINDI) are supported by the National Medical Research Council (NMRC), Singapore (grants 0796/2003, 1176/2008, 1149/2008, STaR/0003/2008, 1249/2010, CG/SERI/2010, CIRG/1371/2013, and CIRG/1417/2015), and Biomedical Research Council (BMRC), Singapore (08/1/35/19/550 and 09/1/35/19/616).

The Singapore Prospective Study Program (SP2) and Living Biobank studies were supported by the individual research grant and clinician scientist award schemes from the National Medical Research Council (NMRC) and the Biomedical Research Council (BMRC) of Singapore, grants from the Singapore Ministry of Health, National University of Singapore and National University Health System, Singapore. Genome Institute of Singapore provided services for genotyping. Genotyping and whole-exome sequencing for Living Biobank was jointly funded the Agency for Science, Technology and Research, Singapore (<https://www.a-star.edu.sg/>) and Merck Sharp & Dohme Corp., Whitehouse Station, NJ USA (<http://www.merck.com>). For the studies above: We thank the study participants and staff who made the research possible, and the Genome Institute of Singapore for genotyping collaboration.

SORBS

This work was supported by grants from the Deutsche Forschungsgemeinschaft (DFG, German Research Foundation – Projektnummer 209933838 – SFB 1052; B03, C01; SPP 1629 TO 718/2- 1). We thank all those who participated in the study. Sincere thanks are given to Dr. Knut Krohn (University of Leipzig) for the genotyping support.

Shanghai Women Health Study

The generation and management of GWAS genotype data for the SWHS was supported by UM1CA182910 and R01CA148677

Special Turku Coronary Risk Factor Intervention Project – Parents

STRIP has been financially supported by Academy of Finland (grants 206374, 251360 and 276861); Juho Vainio Foundation; Finnish Cardiac Research Foundation; Finnish Cultural Foundation; Finnish Ministry of Education and Culture; Sigrid Juselius Foundation; Yrjö Jahnsson Foundation; C.G. Sundell Foundation; Special Governmental Grants for Health Sciences Research, Turku University Hospital; Foundation for Pediatric Research; and Turku University Foundation.

The Hellenic study of Interactions between Single nucleotide polymorphisms and Eating in Atherosclerosis Susceptibility

This work was supported by the British Heart Foundation (BHF) grant RG/14/5/30893 (P.D.) and forms part of the research themes contributing to the translational research portfolios of the Barts Biomedical Research Centre funded by the UK National Institute for Health Research (NIHR)

Taiwan US Diabetic Retinopathy Study

This study was supported by the National Eye Institute of the National Institutes of Health (EY014684) and ARRA Supplement (EY014684-03S1, -04S1), the National Institute of Diabetes and Digestive and Kidney Disease grant DK063491 to the Southern California Diabetes Endocrinology Research Center, the Eye Birth Defects Foundation Inc., the National Science Council, Taiwan (NSC 98-2314-B-075A-002-MY3) and the Taichung Veterans General Hospital, Taichung, Taiwan (TCVGH-1003001C).

Taiwan Hypertension study of Rare Variants

The Rare Variants for Hypertension in Taiwan Chinese (THRV) is supported by the National Heart, Lung, and Blood Institute (NHLBI) grant (R01HL111249) and its participation in TOPMed is supported by an NHLBI supplement (R01HL111249-04S1). THRV is a collaborative study between Washington University in St. Louis, LA BioMed at Harbor UCLA, University of Texas in Houston, Taichung Veterans General Hospital, Taipei Veterans General Hospital, Tri-Service General Hospital, National Health Research Institutes, National Taiwan University, and Baylor University. THRV is based (substantially) on the parent SAPPHIRe study, along with additional population-based and hospital-based cohorts. SAPPHIRe was supported by NHLBI grants (U01HL54527, U01HL54498) and Taiwan funds, and the other cohorts were supported by Taiwan funds.

Tracking Adolescents’ Individual Lives Survey - Population Cohort

TRAILS (TRacking Adolescents’ Individual Lives Survey) is a collaborative project involving various departments of the University Medical Center and University of Groningen, the University of Utrecht, the Radboud Medical Center Nijmegen, and the Parnassia Group, all in the Netherlands. TRAILS has been financially supported by various grants from the Netherlands Organization for Scientific Research NWO (Medical Research Council program grant GB-MW 940-38-011; ZonMW Brainpower grant 100-001-004; ZonMw Risk Behavior and Dependence grants 60-60600-97-118; ZonMw Culture and Health grant 261-98-710; Social Sciences Council medium-sized investment grants GB-MaGW 480-01-006 and GB-MaGW 480-07-001; Social Sciences Council project grants GB-MaGW 452-04-314 and GB-MaGW 452-06-004; NWO large-sized investment grant 175.010.2003.005; NWO Longitudinal Survey and Panel Funding 481-08-013 and 481-11-001; NWO Vici 016.130.002 and 453-16-007/2735; NWO Gravitation 024.001.003), the Dutch Ministry of Justice (WODC), the European Science Foundation (EuroSTRESS project FP-006), the European Research Council (ERC-2017-STG-757364 en ERC-CoG-2015-681466), Biobanking and Biomolecular Resources Research Infrastructure BBMRI-NL (CP 32), the Gratama foundation, the Jan Dekker foundation, the participating universities, and Accare Centre for Child and Adolescent Psychiatry. Statistical analyses were carried out on the Genetic Cluster Computer (http://www.geneticcluster.org), which is financially supported by the Netherlands Scientific Organization (NWO 480-05-003) along with a supplement from the Dutch Brain Foundation. We are grateful to everyone who participated in this research or worked on this project to make it possible.

TWINGENE:

TWINGENE is a substudy of the Swedish Twin Registry which is managed by Karolinska Institutet and receives funding through the Swedish Research Council under the grant no 2017-00641.

Twins UK

TwinsUK receives funding from the Wellcome Trust (212904/Z/18/Z), Medical Research Council. TwinsUK and Massimo Mangino are supported by the National Institute for Health Research (NIHR)-funded BioResource, Clinical Research Facility and Biomedical Research Centre based at Guy’s and St Thomas’ NHS Foundation Trust in partnership with King’s College London. Paraskevi Christofidou is supported by SYSCID project (H2020 contract #733100)

The UK Household Longitudinal Study

The UK Household Longitudinal Study was funded by grants from the Economic & Social Research Council (ES/H029745/1) and the Wellcome Trust (WT098051). UKHLS is led by the Institute for Social and Economic Research at the University of Essex and funded by the Economic and Social Research Council. The survey was conducted by NatCen and the genome-wide scan data were analysed and deposited by the Wellcome Trust Sanger Institute. Information on how to access the data can be found on the Understanding Society website (https://www.understandingsociety.ac.uk/).

Vejle Diabetes Biobank

The Vejle Diabetes Biobank was supported by The Danish Research Council for Independent Research and by Region of Southern Denmark.

The Viking Health Study- Shetland

The Viking Health Study – Shetland (VIKING) was supported by the MRC Human Genetics Unit quinquennial programme grant “QTL in Health and Disease”. DNA extractions and genotyping were performed at the Edinburgh Clinical Research Facility, University of Edinburgh. We would like to acknowledge the invaluable contributions of the research nurses in Shetland, the administrative team in Edinburgh and the people of Shetland.

Whitehall II Study

The Whitehall II Study has been supported by grants from the UK Medical Research Council (K013351, R024227, S011676); the British Heart Foundation (PG/11/63/29011 and RG/13/2/30098); the British Health and Safety Executive; the British Department of Health; the National Heart, Lung, and Blood Institute (R01HL036310); the National Institute on Ageing, National Institute of Health (R01AG056477, R01AG034454); and the Economic and Social Research Council (ES/J023299/1).

Women's Health Initiative

The WHI program is funded by the National Heart, Lung, and Blood Institute, National Institutes of Health, U.S. Department of Health and Human Services through contracts HHSN268201600018C, HHSN268201600001C, HHSN268201600002C, HHSN268201600003C, and HHSN268201600004C. We also acknowledge the contributions of the following individuals:

Program Office: (National Heart, Lung, and Blood Institute, Bethesda, Maryland) Jacques Rossouw, Shari Ludlam, Joan McGowan, Leslie Ford, and Nancy Geller

Clinical Coordinating Center: (Fred Hutchinson Cancer Research Center, Seattle, WA) Garnet Anderson, Ross Prentice, Andrea LaCroix, and Charles Kooperberg

Investigators and Academic Centers: (Brigham and Women's Hospital, Harvard Medical School, Boston, MA) JoAnn E. Manson; (MedStar Health Research Institute/Howard University, Washington, DC) Barbara V. Howard; (Stanford Prevention Research Center, Stanford, CA) Marcia L. Stefanick; (The Ohio State University, Columbus, OH) Rebecca Jackson; (University of Arizona, Tucson/Phoenix, AZ) Cynthia A. Thomson; (University at Buffalo, Buffalo, NY) Jean Wactawski-Wende; (University of Florida, Gainesville/Jacksonville, FL) Marian Limacher; (University of Iowa, Iowa City/Davenport, IA) Jennifer Robinson; (University of Pittsburgh, Pittsburgh, PA) Lewis Kuller; (Wake Forest University School of Medicine, Winston-Salem, NC) Sally Shumaker; (University of Nevada, Reno, NV) Robert Brunner

Women’s Health Initiative Memory Study: (Wake Forest University School of Medicine, Winston-Salem, NC) Mark Espeland

Women's Genome Health Study

The WGHS is supported by the National Heart, Lung, and Blood Institute (HL043851 and HL080467) and the National Cancer Institute (CA047988 and UM1CA182913). Additional support and funding for genotyping was provided by Amgen.

**Consortium Authors: VA Million Veteran Program**

MVP Executive Committee

-	Co-Chair: J. Michael Gaziano, M.D., M.P.H.

VA Boston Healthcare System, 150 S. Huntington Avenue, Boston, MA 02130

-	Co-Chair: Sumitra Muralidhar, Ph.D.

US Department of Veterans Affairs, 810 Vermont Avenue NW, Washington, DC 20420

-	Rachel Ramoni, D.M.D., Sc.D., Chief VA Research and Development Officer

US Department of Veterans Affairs, 810 Vermont Avenue NW, Washington, DC 20420

-	Jean Beckham, Ph.D.

Durham VA Medical Center, 508 Fulton Street, Durham, NC 27705

-	Kyong-Mi Chang, M.D.

Philadelphia VA Medical Center, 3900 Woodland Avenue, Philadelphia, PA 19104

-	Christopher J. O’Donnell, M.D., M.P.H.

VA Boston Healthcare System, 150 S. Huntington Avenue, Boston, MA 02130

-	Philip S. Tsao, Ph.D.

VA Palo Alto Health Care System, 3801 Miranda Avenue, Palo Alto, CA 94304

-	James Breeling, M.D., Ex-Officio

US Department of Veterans Affairs, 810 Vermont Avenue NW, Washington, DC 20420

-	Grant Huang, Ph.D., Ex-Officio

US Department of Veterans Affairs, 810 Vermont Avenue NW, Washington, DC 20420

-	JP Casas Romero, M.D., Ph.D., Ex-Officio

VA Boston Healthcare System, 150 S. Huntington Avenue, Boston, MA 02130

MVP Program Office

-	Sumitra Muralidhar, Ph.D.

US Department of Veterans Affairs, 810 Vermont Avenue NW, Washington, DC 20420

-	Jennifer Moser, Ph.D.

US Department of Veterans Affairs, 810 Vermont Avenue NW, Washington, DC 20420

MVP Recruitment/Enrollment

-	Recruitment/Enrollment Director/Deputy Director, Boston – Stacey B. Whitbourne, Ph.D.; Jessica V. Brewer, M.P.H.

VA Boston Healthcare System, 150 S. Huntington Avenue, Boston, MA 02130

-	MVP Coordinating Centers

o	Clinical Epidemiology Research Center (CERC), West Haven – Mihaela Aslan, Ph.D.

West Haven VA Medical Center, 950 Campbell Avenue, West Haven, CT 06516

o	Cooperative Studies Program Clinical Research Pharmacy Coordinating Center, Albuquerque – Todd Connor, Pharm.D.; Dean P. Argyres, B.S., M.S.

New Mexico VA Health Care System, 1501 San Pedro Drive SE, Albuquerque, NM 87108

o	Genomics Coordinating Center, Palo Alto – Philip S. Tsao, Ph.D.

VA Palo Alto Health Care System, 3801 Miranda Avenue, Palo Alto, CA 94304

o	MVP Boston Coordinating Center, Boston - J. Michael Gaziano, M.D., M.P.H.

VA Boston Healthcare System, 150 S. Huntington Avenue, Boston, MA 02130

o	MVP Information Center, Canandaigua – Brady Stephens, M.S.

Canandaigua VA Medical Center, 400 Fort Hill Avenue, Canandaigua, NY 14424

-	VA Central Biorepository, Boston – Mary T. Brophy M.D., M.P.H.; Donald E. Humphries, Ph.D.; Luis E. Selva, Ph.D.

VA Boston Healthcare System, 150 S. Huntington Avenue, Boston, MA 02130

-	MVP Informatics, Boston – Nhan Do, M.D.; Shahpoor Shayan

VA Boston Healthcare System, 150 S. Huntington Avenue, Boston, MA 02130

-	MVP Data Operations/Analytics, Boston – Kelly Cho, Ph.D.

VA Boston Healthcare System, 150 S. Huntington Avenue, Boston, MA 02130

MVP Science

-	Science Operations – Christopher J. O’Donnell, M.D., M.P.H.

VA Boston Healthcare System, 150 S. Huntington Avenue, Boston, MA 02130

-	Genomics Core - Christopher J. O’Donnell, M.D., M.P.H.; Saiju Pyarajan Ph.D.

VA Boston Healthcare System, 150 S. Huntington Avenue, Boston, MA 02130

Philip S. Tsao, Ph.D.

VA Palo Alto Health Care System, 3801 Miranda Avenue, Palo Alto, CA 94304

-	Phenomics Core- Kelly Cho, M.P.H, Ph.D.

VA Boston Healthcare System, 150 S. Huntington Avenue, Boston, MA 02130

-	Data and Computational Sciences – Saiju Pyarajan, Ph.D.

VA Boston Healthcare System, 150 S. Huntington Avenue, Boston, MA 02130

-	Statistical Genetics – Elizabeth Hauser, Ph.D.

Durham VA Medical Center, 508 Fulton Street, Durham, NC 27705

Yan Sun, Ph.D.

Atlanta VA Medical Center, 1670 Clairmont Road, Decatur, GA 30033

Hongyu Zhao, Ph.D.

West Haven VA Medical Center, 950 Campbell Avenue, West Haven, CT 06516

Current MVP Local Site Investigators

-	Atlanta VA Medical Center (Peter Wilson, M.D.)

1670 Clairmont Road, Decatur, GA 30033

-	Bay Pines VA Healthcare System (Rachel McArdle, Ph.D.)

10,000 Bay Pines Blvd Bay Pines, FL 33744

-	Birmingham VA Medical Center (Louis Dellitalia, M.D.)

700 S. 19th Street, Birmingham AL 35233

-	Central Western Massachusetts Healthcare System (Kristin Mattocks, Ph.D., M.P.H.)

421 North Main Street, Leeds, MA 01053

-	Cincinnati VA Medical Center (John Harley, M.D., Ph.D.)

3200 Vine Street, Cincinnati, OH 45220

-	Clement J. Zablocki VA Medical Center (Jeffrey Whittle, M.D., M.P.H.)

5000 West National Avenue, Milwaukee, WI 53295

-	VA Northeast Ohio Healthcare System (Frank Jacono, M.D.)

10701 East Boulevard, Cleveland, OH 44106

-	Durham VA Medical Center (Jean Beckham, Ph.D.)

508 Fulton Street, Durham, NC 27705

-	Edith Nourse Rogers Memorial Veterans Hospital (John Wells., Ph.D.)

200 Springs Road, Bedford, MA 01730

-	Edward Hines, Jr. VA Medical Center (Salvador Gutierrez, M.D.)

5000 South 5th Avenue, Hines, IL 60141

-	Veterans Health Care System of the Ozarks (Gretchen Gibson, D.D.S., M.P.H.)

1100 North College Avenue, Fayetteville, AR 72703

-	Fargo VA Health Care System (Kimberly Hammer, Ph.D.)

2101 N. Elm, Fargo, ND 58102

-	VA Health Care Upstate New York (Laurence Kaminsky, Ph.D.)

113 Holland Avenue, Albany, NY 12208

-	New Mexico VA Health Care System (Gerardo Villareal, M.D.)

1501 San Pedro Drive, S.E. Albuquerque, NM 87108

-	VA Boston Healthcare System (Scott Kinlay, M.B.B.S., Ph.D.)

150 S. Huntington Avenue, Boston, MA 02130

-	VA Western New York Healthcare System (Junzhe Xu, M.D.)

3495 Bailey Avenue, Buffalo, NY 14215-1199

-	Ralph H. Johnson VA Medical Center (Mark Hamner, M.D.)

109 Bee Street, Mental Health Research, Charleston, SC 29401

-	Columbia VA Health Care System (Roy Mathew, M.D.)

6439 Garners Ferry Road, Columbia, SC 29209

-	VA North Texas Health Care System (Sujata Bhushan, M.D.)

4500 S. Lancaster Road, Dallas, TX 75216

-	Hampton VA Medical Center (Pran Iruvanti, D.O., Ph.D.)

100 Emancipation Drive, Hampton, VA 23667

-	Richmond VA Medical Center (Michael Godschalk, M.D.)

1201 Broad Rock Blvd., Richmond, VA 23249

-	Iowa City VA Health Care System (Zuhair Ballas, M.D.)

601 Highway 6 West, Iowa City, IA 52246-2208

-	Eastern Oklahoma VA Health Care System (Douglas Ivins, M.D.)

1011 Honor Heights Drive, Muskogee, OK 74401

-	James A. Haley Veterans’ Hospital (Stephen Mastorides, M.D.)

13000 Bruce B. Downs Blvd, Tampa, FL 33612

-	James H. Quillen VA Medical Center (Jonathan Moorman, M.D., Ph.D.)

Corner of Lamont & Veterans Way, Mountain Home, TN 37684

-	John D. Dingell VA Medical Center (Saib Gappy, M.D.)

4646 John R Street, Detroit, MI 48201

-	Louisville VA Medical Center (Jon Klein, M.D., Ph.D.)

800 Zorn Avenue, Louisville, KY 40206

-	Manchester VA Medical Center (Nora Ratcliffe, M.D.)

718 Smyth Road, Manchester, NH 03104

-	Miami VA Health Care System (Hermes Florez, M.D., Ph.D.)

1201 NW 16th Street, 11 GRC, Miami FL 33125

-	Michael E. DeBakey VA Medical Center (Olaoluwa Okusaga, M.D.)

2002 Holcombe Blvd, Houston, TX 77030

-	Minneapolis VA Health Care System (Maureen Murdoch, M.D., M.P.H.)

One Veterans Drive, Minneapolis, MN 55417

-	N. FL/S. GA Veterans Health System (Peruvemba Sriram, M.D.)

1601 SW Archer Road, Gainesville, FL 32608

-	Northport VA Medical Center (Shing Shing Yeh, Ph.D., M.D.)

79 Middleville Road, Northport, NY 11768

-	Overton Brooks VA Medical Center (Neeraj Tandon, M.D.)

510 East Stoner Ave, Shreveport, LA 71101

-	Philadelphia VA Medical Center (Darshana Jhala, M.D.)

3900 Woodland Avenue, Philadelphia, PA 19104

-	Phoenix VA Health Care System (Samuel Aguayo, M.D.)

650 E. Indian School Road, Phoenix, AZ 85012

-	Portland VA Medical Center (David Cohen, M.D.)

3710 SW U.S. Veterans Hospital Road, Portland, OR 97239

-	Providence VA Medical Center (Satish Sharma, M.D.)

830 Chalkstone Avenue, Providence, RI 02908

-	Richard Roudebush VA Medical Center (Suthat Liangpunsakul, M.D., M.P.H.)

1481 West 10th Street, Indianapolis, IN 46202

-	Salem VA Medical Center (Kris Ann Oursler, M.D.)

1970 Roanoke Blvd, Salem, VA 24153

-	San Francisco VA Health Care System (Mary Whooley, M.D.)

4150 Clement Street, San Francisco, CA 94121

-	South Texas Veterans Health Care System (Sunil Ahuja, M.D.)

7400 Merton Minter Boulevard, San Antonio, TX 78229

-	Southeast Louisiana Veterans Health Care System (Joseph Constans, Ph.D.)

2400 Canal Street, New Orleans, LA 70119

-	Southern Arizona VA Health Care System (Paul Meyer, M.D., Ph.D.)

3601 S 6th Avenue, Tucson, AZ 85723

-	Sioux Falls VA Health Care System (Jennifer Greco, M.D.)

2501 W 22nd Street, Sioux Falls, SD 57105

-	St. Louis VA Health Care System (Michael Rauchman, M.D.)

915 North Grand Blvd, St. Louis, MO 63106

-	Syracuse VA Medical Center (Richard Servatius, Ph.D.)

800 Irving Avenue, Syracuse, NY 13210

-	VA Eastern Kansas Health Care System (Melinda Gaddy, Ph.D.)

4101 S 4th Street Trafficway, Leavenworth, KS 66048

-	VA Greater Los Angeles Health Care System (Agnes Wallbom, M.D., M.S.)

11301 Wilshire Blvd, Los Angeles, CA 90073

-	VA Long Beach Healthcare System (Timothy Morgan, M.D.)

5901 East 7th Street Long Beach, CA 90822

-	VA Maine Healthcare System (Todd Stapley, D.O.)

1 VA Center, Augusta, ME 04330

-	VA New York Harbor Healthcare System (Scott Sherman, M.D., M.P.H.)

423 East 23rd Street, New York, NY 10010

-	VA Pacific Islands Health Care System (George Ross, M.D.)

459 Patterson Rd, Honolulu, HI 96819

-	VA Palo Alto Health Care System (Philip Tsao, Ph.D.)

3801 Miranda Avenue, Palo Alto, CA 94304-1290

-	VA Pittsburgh Health Care System (Patrick Strollo, Jr., M.D.)

University Drive, Pittsburgh, PA 15240

-	VA Puget Sound Health Care System (Edward Boyko, M.D.)

1660 S. Columbian Way, Seattle, WA 98108-1597

-	VA Salt Lake City Health Care System (Laurence Meyer, M.D., Ph.D.)

500 Foothill Drive, Salt Lake City, UT 84148

-	VA San Diego Healthcare System (Samir Gupta, M.D., M.S.C.S.)

3350 La Jolla Village Drive, San Diego, CA 92161

-	VA Sierra Nevada Health Care System (Mostaqul Huq, Pharm.D., Ph.D.)

975 Kirman Avenue, Reno, NV 89502

-	VA Southern Nevada Healthcare System (Joseph Fayad, M.D.)

6900 North Pecos Road, North Las Vegas, NV 89086

-	VA Tennessee Valley Healthcare System (Adriana Hung, M.D., M.P.H.)

1310 24th Avenue, South Nashville, TN 37212

-	Washington DC VA Medical Center (Jack Lichy, M.D., Ph.D.)

50 Irving St, Washington, D. C. 20422

-	W.G. (Bill) Hefner VA Medical Center (Robin Hurley, M.D.)

1601 Brenner Ave, Salisbury, NC 28144

-	White River Junction VA Medical Center (Brooks Robey, M.D.)

163 Veterans Drive, White River Junction, VT 05009

-	William S. Middleton Memorial Veterans Hospital (Robert Striker, M.D., Ph.D.)

2500 Overlook Terrace, Madison, WI 53705
